# “Decoding Mobile Genetic Elements, Virulence Factors, and Antibiotic Resistance Genes Driving Biofilm Formation in MRSA via Network Analysis”

**DOI:** 10.1101/2025.01.30.635750

**Authors:** Daraksha Iram, Manish Singh Sansi, Indrajit Ganguly, Sanjeev Singh, Satpal Dixit

**Author notes:** **Corresponding authors:** Satpal Dixit-, Daraksha Iram. Daraksha Iram and Manish Singh Sansi contributed equally to this article.

## Abstract

This study aimed to analyze the genome of the Methicillin-resistant *Staphylococcus aureus* (MRSA) strain d1418m22 through whole-genome sequencing (WGS) and comprehensive bioinformatics analysis with six references strains. *De novo* assembly resulted in 61 scaffolds with a total genome size of 2.78 Mb. Functional annotation revealed 2,625 predicted genes, including those involved in metabolism, cellular processes, and virulence. Comparative genomic analysis identified 20 antibiotic resistance genes, including those conferring resistance to beta-lactams, fluoroquinolones, and aminoglycosides. In addition, 21 virulence factors (VFs) were identified, including Panton-Valentine leukocidin (PVL) and various enterotoxins. The presence of staphylococcal cassette chromosome mec (*SCCmec*) type IVa and mobile genetic elements (MGEs), such as prophages and transposons, underscored the role of horizontal gene transfer (HGT) in the evolution and transmission of resistance and virulence factors. Interaction network analysis identified key hub genes involved in various cellular processes, including biofilm formation. This comprehensive genomic analysis provides valuable insights into the genetic makeup of MRSA strain d1418m22, contributing to a better understanding of its pathogenicity and potential public health implications. Six reference MRSA strains were isolated from (MRSA-AMRF4 and MRSA-AMRF5) eye infections, (MRSA and MRSA-15) wound pus, (VMRSA-WC071, and VMRSA-WC081) urine samples. The analysis identified resistance genes, virulence factors, GC content, ANI values, and *SCCmec* elements, which were found to be similar to those present in the resistant MRSA d1418m22 strain genome.

## 1.0. Introduction

*Staphylococcus* species can cause a wide range of infections, from acute to chronic, leading to significant morbidity in both humans and animals (Pattabhiramaiah and Mallikarjunaiah, 2023). While these bacteria are commonly commensals in humans, they can transition into pathogenic forms (Becker et al., 2014). *Staphylococcus aureus* and *Staphylococcus haemolyticus* are clinically significant bacterial species. *S. aureus* is recognized as a major human pathogen, associated with a wide spectrum of infections, ranging from superficial skin infections to life-threatening conditions like septicemia, pneumonia, and endocarditis (Tong et al., 2015). The emergence of antibiotic-resistant strains of *S. aureus* poses a significant challenge to public health, both in community and healthcare settings (Stefani et al., 2012; Chambers et al., 2009). *S. haemolyticus* is frequently isolated in bloodstream infections. *S. aureus* is a significant pathogen in bovine mastitis. It has the ability to adhere to and invade mammary epithelial cells, leading to the establishment of chronic intramammary infections (IMIs) (Maity et al., 2020). Surgical site infections (SSIs) represent a significant healthcare burden, affecting 2-5% of patients undergoing surgical procedures worldwide (Smyth et al., 2008; Magill et al., 2012). These infections account for 14-33% of all healthcare-associated infections (Smyth et al., 2008). *S. aureus* is a leading cause of SSIs (Dhar et al., 2014; Mpogoro et al., 2014).

The *Staphylococcus* genus includes 53 species and 28 subspecies, with *S. aureus* being a primary cause of chronic IMIs (Vanderhaeghen et al., 2015). Despite extensive research, the molecular mechanisms of *S. aureus* in IMIs are not fully understood. Typically, bacteria detect host signals and regulate gene expression to adapt and establish infections (Elhawy et al., 2021). *S. aureus* has several VFs that help it adhere, invade, and evade the immune system, and these are encoded either in the bacterial genome or on MGEs (Naushad et al., 2019). Molecular typing methods are crucial for characterizing the genetic diversity of *S. aureus*. Multilocus sequence typing (MLST) is a widely used technique for identifying and tracking the dissemination of different *S. aureus* clones (Lindsay et al., 2014). Spa typing complements MLST by providing high-resolution data on the genetic evolution of *S. aureus* strains. Furthermore, spa typing offers valuable insights into the role of protein A in *S. aureus* pathogenesis (Pobiega et al., 2009). Characterizing the *SCCmec* elements, particularly the presence of the mecA gene, is crucial for understanding methicillin resistance in *Staphylococcus aureus* (MRSA) (Funaki et al., 2019; Sabri et al., 2013). Molecular epidemiology of *Staphylococcus species* involved in intramammary infections (IMIs) in dairy cattle has been extensively studied. Various typing techniques, including multi-locus enzyme electrophoresis (MLEE), pulsed-field gel electrophoresis (PFGE), multi-locus sequence typing (MLST), and multiple-locus variable number tandem repeat analysis (MLVA), have been employed to investigate the genetic diversity and transmission of these pathogens. Additionally, techniques such as random amplification of polymorphic DNA (RAPD) and spa typing have been utilized to analyze genetic diversity and track epidemiological patterns within *S. aureus* populations associated with IMIs (Li, 2008).

*Staphylococci* may act as reservoirs for antibiotic-resistance-genes, which can potentially be transferred to other species, including *S. aureus* (Froggatt et al., 1989; Maarouf et al., 2020). Effective infection control depends on thorough epidemiological monitoring and genetic profiling (Steinig et al., 2019; Pobiega et al., 2013). *S. aureus* pathogenesis begins with teat colonization, progressing into the intramammary space through gradual colonization or pressure fluctuations from milking equipment (Maity et al., 2020; Vargová et al., 2023). Methicillin-resistant *S. aureus* (MRSA) complicates treatment due to multi-drug resistance, while rising antibiotic resistance worsens mastitis management, especially with widespread antibiotic use in dairy farming (Klibi et al., 2018). Practices like dry cow therapy promote the development of multi-drug-resistant *S. aureus* strains (Liu et al., 2020; Omwenga et al., 2021; Gelalcha et al., 2022).

Whole-genome sequencing (WGS) is a powerful tool for typing, differentiating, and clustering *staphylococcal* isolates, making it vital for outbreak control (Harris et al., 2013; Eyre et al., 2012). It enables the identification of outbreak clones or clades—groups of related isolates with shared characteristics likely derived from a common ancestor (Struelens et al., 1996; Besser et al., 2019). WGS can fill gaps in staphylococcal management, particularly in regions with limited surveillance, such as Egypt and other non-European Mediterranean countries, where staphylococcal epidemiology is underexplored (Asadollahi et al., 2021; Tokajian et al., 2014). While traditional genetic methods may lack the sensitivity to detect subtle genetic differences, WGS offers higher resolution for studying microevolution, phylogenetic relationships, and inter-species variations (Sivakumar et al., 2023).

In the current study, we used whole-genome sequencing (WGS) to investigate the genomic diversity, VFs, and antimicrobial resistance mechanisms in the MRSA strain d1418m22 genome, isolated from knee wound pus samples collected in a hospital. This isolate was identified as gram-positive cocci and catalase-positive. This study explores the antimicrobial resistance patterns and molecular characteristics of *Staphylococcus aureus* MRSA. Specifically, it focuses on analyzing the whole genome sequence (WGS) of the MRSA strain d1418m22. The study further evaluates the genomic diversity and provides a genotypic prediction of antimicrobial resistance in MRSA isolated from a patient in India.

## 2.0. Material method

### 2.1. Bacterial Strain MRSA genome collection

The MRSA strain d1418m22, featuring a 2.78-MB *de novo* assembled whole-genome sequence, was obtained from the NCBI genome database. The associated details are as follows: accession number JAPDDY000000000, BioProject ID PRJNA894174, BioSample ID SAMN31439881, and SRA project number SRR24389989. The de novo assembly was performed using KmerGenie (Chikhi et al., 2014), Velvet (Zerbino et al., 2008), SPAdes (Bankevich et al., 2012), and GapFiller (Nadalin et al., 2012). Default settings were applied for all software. The analysis was conducted by comparing d1418m22 with six assembled strains available in the NCBI Genome database. The reference strains included two isolates from eye infection samples (NCBI GenBank IDs: <u>GCA_015219905.1</u> and <u>GCA_015219885.1</u>), two isolates from wound pus samples (<u>GCA_033738955.1</u> and <u>GCA_000695215.1</u>), and two isolates from urine samples (<u>GCA_022693325.1</u> and <u>GCA_022693305.1</u>).

### 2.2. Computational Analysis of MRSA Genome

Antimicrobial resistance (AMR) genes were identified using the Comprehensive Antibiotic Resistance Database (CARD) (Alcock et al., 2023).

CARD provided insights into the broad mechanisms of antibiotic-resistance, the associated drug classes, and, in specific cases, the particular antibiotics to which resistance is conferred. Additional bioinformatics tools included ResFinder (v4.1.0) (Zankari et al., 2017; Bortolaia et al., 2020), available at (https://cge.food.dtu.dk/services/ResFinder/), for precise antimicrobial resistance gene assignment. While Plasmid Finder (v2.1.0) (Carattoli et al., 2014) was used to detect plasmid-encoded replicases, contributing to understanding MGEs. These tools were accessed via the Center for Genomic-Epidemiology platform (https://cge.cbs.dtu.dk/services/). To visualize the analyzed genome, a circular genome map was generated using CGView.js through the Proksee platform (Grant et al., 2008). This visualization facilitated an integrated view of AMR genes, virulence factors, and plasmid elements within the genomic context, providing a comprehensive understanding of the genomic architecture and its implications for antimicrobial resistance.

### 2.3. Identification and assignment of Virulence Factors and Staphylococcal Chromosomal Cassette mec (SCCmec) Elements

VFs were identified using VFanalyzer, an analytical tool available in the Virulence Factors-Database (VFDB) (http://www.mgc.ac.cn/VFs/main.htm, accessed on 25 November 2024) (Liu et al., 2019; Chen et al., 2012). The characterization of Staphylococcal Chromosomal Cassette mec (SCCmec) elements was performed using SCCmec Finder 1.2, hosted by the Center for Genomic Epidemiology (https://cge.food.dtu.dk/services/SCCmecFinder/, accessed on 1 December 2024) (Carattoli et al., 2020; Carattoli et al., 2014), as well as resources from the International Working Group on Staphylococcal Cassette Chromosome (IWC-SCC) (https://www.sccmec.org/index.php/en/, accessed on 1 December 2024). To identify potential prophages within the genome sequence, the Phage-Search-Tool-Enhanced Release (PHASTER) (https://phaster.ca/, accessed on 1 December 2022) was employed (Zhou et al., 2011; Arndt et al., 2016). All these bioinformatics tools were accessed via the Center for Genomic-Epidemiology’s integrated platform (https://cge.cbs.dtu.dk/services/). The resulting genomic data were visualized as circular genome maps using CGView.js, available through Proksee (https://proksee.ca/, accessed on 20 November 2024) (Grant et al., 2008). This comprehensive approach ensured detailed and accurate analysis of the genome for virulence factors, SCCmec elements, prophages, and plasmid-encoded elements. The genome analysis employed advanced bioinformatics tools for comprehensive characterization and functional annotation. CRISPRCas Finder 4.2.20 was utilized to identify and analyze CRISPR-Cas systems, shedding light on bacterial adaptive immunity. Proksee (Tool Version 1.0) was applied to detect open reading frames (ORFs), crucial for predicting protein-coding regions. Genome annotation was conducted using Prokka 1.14.6 (Seemann et al., 2014), automating the identification of genes, coding sequences, rRNAs, tRNAs, and other genomic features, ensuring detailed and high-quality annotations. To predict HGT events, by alien hunter 1.7 (Vernikos et al., 2006) identified genomic regions likely acquired through HGT, highlighting foreign DNA integration’s role in bacterial evolution. Additionally, mobileOG-db (beatrix-1.6) (Brown et al., 2022), characterized MGEs, including plasmids and transposons, critical for understanding genome dynamics and the dissemination of antibiotic resistance genes.

### 2.4. Comparison analysis BLAST of genome sequence

The *Staphylococcus aureus* MRSA strain d1418m22 was analyzed and compared to other resistant reference strains using the BLAST+ 2.15.0 tool. Additionally, the FastANI 1.34 tool was employed for rapid, alignment-free computation of whole-genome Average Nucleotide Identity (ANI).

### 2.5. STRING network identified of top resistant and biofilm formation genes

The top antibiotic resistance and biofilm formation genes were identified from genome annotation results and analyzed using the STRING-database (https://string-db.org/) to construct a protein-protein interaction (PPI)-network (Viering et al., 2022). STRING parameters, including a medium confidence score (≥ 0.4), were applied, and the resulting interaction data was exported for visualization in Cystoscape (v3.9.0) (Shannon et al., 2003). The PPI network was imported into Cystoscape, where node attributes, such as gene names and functional categories, were mapped. Degree centrality, calculated using the “Network Analyzer” plugin, identified hub genes with the most interactions, reflecting their critical roles (Chin et al., 2014). Sub networks of hub genes and immediate neighbors were extracted for detailed analysis. This integrated approach highlighted key genes involved in resistance and biofilm formation, offering potential targets for therapeutic intervention and advancing understanding of microbial mechanisms. The full networks are available for interactive visualization in Cystoscape (Shannon et al., 2003). The methodology offers significant insights into genetic elements that drive bacterial survival, pathogenicity, and resistance mechanisms.

## 3.0. Result and discussion

### 3.1. Genome project history

The genome sequence of the MRSA strain d1418m22 has been deposited in the GenBank WGS database. A summary of the results is provided in Table 1.

### 3.2. Genomic features of bacterial strain

The data in Table 1 indicates that the draft genome sequence of MRSA strain d1418m22 comprises a total genome size of 2,784,608 bp (excluding gaps). The assembly features 2,583 predicted coding sequences (CDS), 55 tRNAs, and 6 rRNAs. The N50 scaffold length is 130,908 bp, with the largest assembled contigs measuring 290,662 bp. The libraries were sequenced using the Illumina-NovaSeq 6000 platform (2 × 150 bp chemistry), generating 3–4 GB of data per sample. The average library size was 429 bp for MRSA strain d1418m22 genome. *De-novo* assembly of reads was performed using SPAdes 3.15.4, with additional tools including KmerGenie, Velvet, and GapFiller employed to refine the assemblies (Iram et al., 2024). The final draft genomes were determined to be 100% complete using BUSCO (version 5.4.6). Default settings were applied across all software tools to ensure consistency in the assembly process. The present study identified 20,229,752 high-quality reads with a guanine-cytosine content of 32.72%. De novo assembly produced 61 scaffolds, yielding an N50 of 130,908 bp, a maximum scaffold size of 290,662 bp, and a genome size of 2.78 Mb sequenced MRSA strains (Iram et al., 2024). The average scaffold length was 45,649 bp, with scaffold sizes ranging from a maximum of 290,662 bp to a minimum of 548 bp, as detailed in Table 1. Scaffolds were further analyzed and categorized based on gene length distribution, as illustrated in Figure 1a. In the MRSA 1418 strain, the minimum gene length was ≤200 bp (964 genes), while the maximum length exceeded 5,000 bp (1 gene). The top alignment for MRSA closely matched other species within the *Staphylococcus* genus, particularly *Staphylococcus aureus*. This strong genetic similarity highlights the relatedness among *Staphylococcus* species, with *S. aureus* emerging as the most prominent match, as shown in Figure 1b. A similar study reported the draft genome of *S. aureus* strain SO-1977 spanning 2,827,644-bp with a GC content of 32.8%. The genome includes 2,629 coding sequences, 51 tRNAs, and 4 rRNAs. The assembly comprises 151-contigs with an N50 of 62,783 bp, and the largest contigs measures 146,886-bp, offering insights into its genetic composition and functional potential (Ali et al., 2019). Forde et al. (2022) sequenced 2,660 isolates over four years, including MDR gram-negative bacilli (CPE: 293, ESBL: 1309), MRSA (620), and VRE (433), resulting in 379 clinical reports. Core genome SNP analysis showed 33% of isolates forming 76 clusters, with 43 confined to target hospitals, indicating intra-hospital transmission. The remaining 33 clusters suggested inter-hospital or community spread. Notably, negligible non-multi-resistant MRSA transmission in one hospital led to changes in infection control policies. These studies underscore the importance of genomic surveillance in understanding antimicrobial resistance and optimizing infection control strategies.

**Figure 1.**
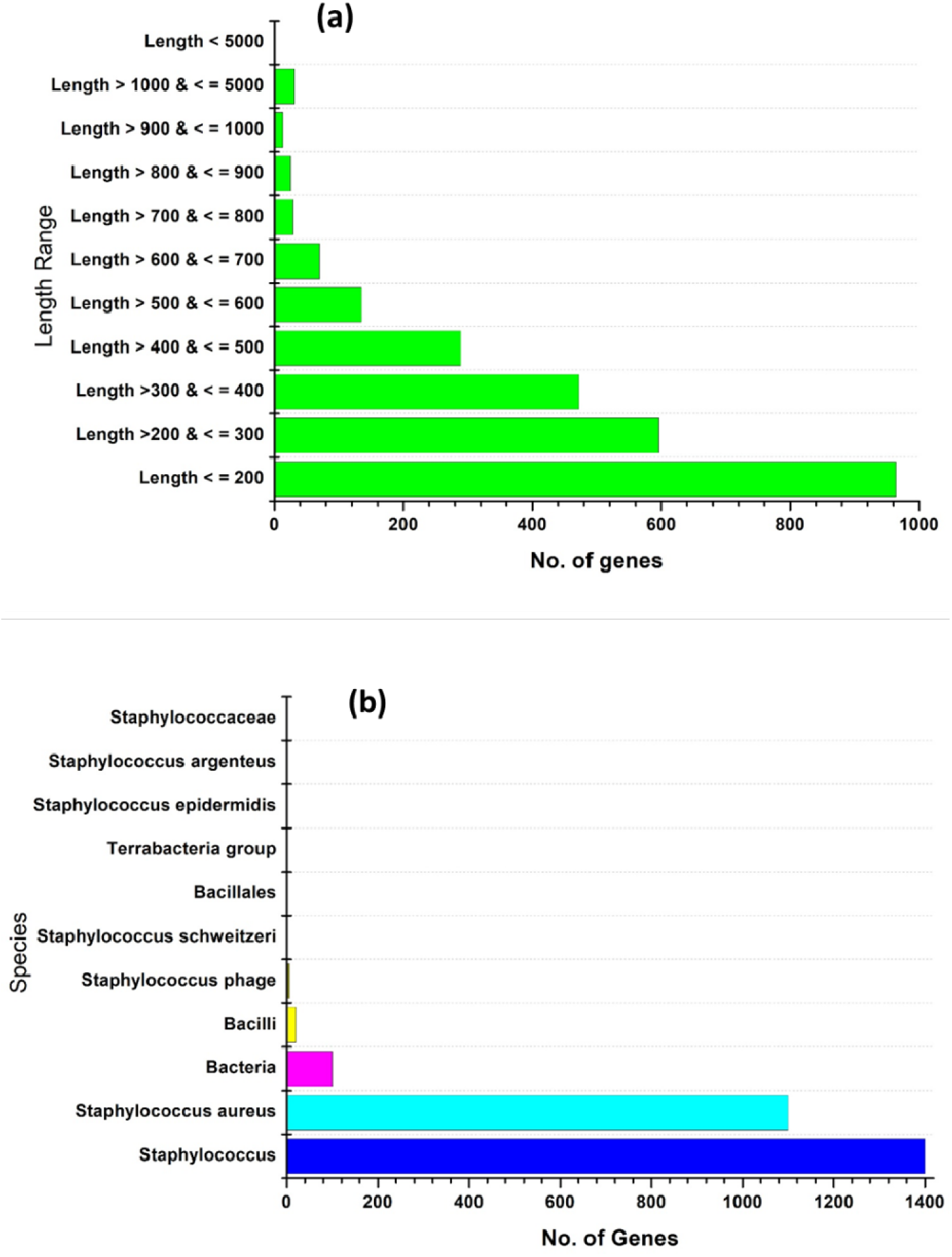
**(a)** Gene length distribution for MRSA strain d1418m22 genome was length-range is plotted on X-axis and its corresponding gene number on Y-axis. **(b)** *S. aureus* top hit with *Staphylococcus species* such as *Staphylococcus aureus*.

### 3.3. Blast statistics against COG, Pfam and Uniprot database

All 2,625 predicted genes from the MRSA strain d1418m22 genome were simultaneously analyzed for similarity search against the Uniprot, COG, and Pfam databases via BLASTP with an e-value threshold of 1e^-5^. The results of the similarity search across these three databases are as follows: In the Uniprot, COG, and Pfam databases, a total of 2,625 proteins were analyzed, with 2,060, 2,130, and 1,770 proteins showing significant hits, respectively. Meanwhile, 565, 495, and 855 proteins, respectively, showed no significant hits in these databases, as illustrated in Figure 2 (a, b, and c). A previous study suggested that the genome of the MRSA strain SO-1977 contained 2,629 predicted coding sequences (CDSs) (Ali et al., 2019).

**Figure 2.**
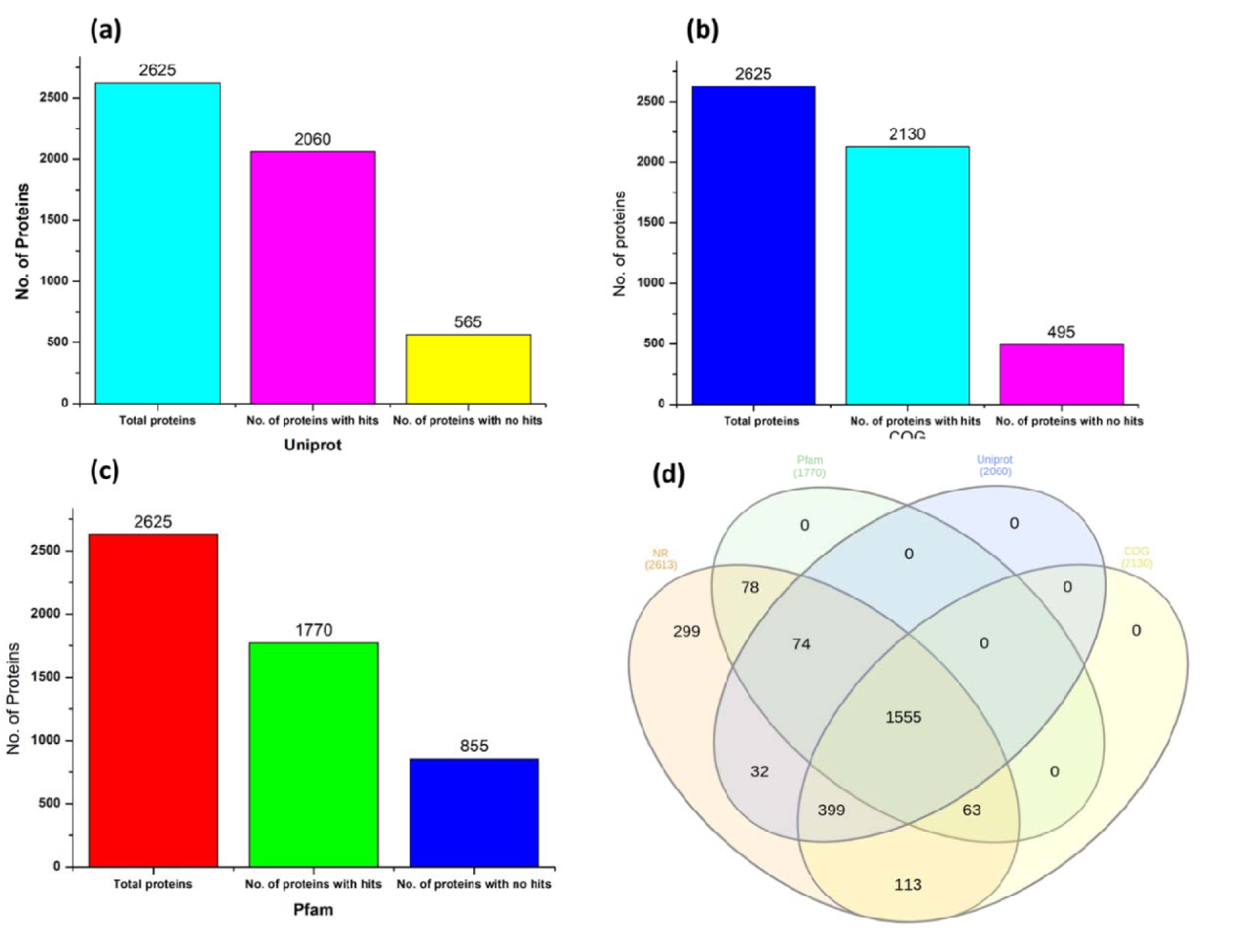
Similarity searched against all three databases are as follows: **(a)** Uniprot **(b)** COG **(c)** Pfam-database using BLASTP with an e-value-threshold of 1e^-5^. Results for similarity search.

### 3.4. Functional distribution counts for Clusters of Orthologous Groups as class wise

The functional categorization of genes in the MRSA strain d1418m22 was performed using the COG (Clusters of Orthologous Groups) classification system. This system organizes genes into specific functional categories, reflecting diverse biological roles, cellular structures, and molecular mechanisms. Bacterial genomic features were further analysed using subsystem technology, which groups genes based on their involvement in particular biological functions within a structural context. Genome analysis of strain d1418m22, conducted using the RAST-subsystem-server, revealed 26 functional categories, as shown in Figure 3. The largest number of genes were found in the amino acid-transport and metabolism category (n = 223), followed by categories such as translation, ribosomal-structure, and biogenesis (n = 217); general function prediction (n = 177); inorganic ion-transport and metabolism (n = 168); carbohydrate-transport and metabolism (n = 162); transcription (n = 155); coenzyme-transport and metabolism (n = 147); cell wall/membrane/envelope biogenesis (n = 146); replication, recombination, and repair (n = 113); energy production and conversion (n = 110); and lipid-transport and metabolism (n = 101). Other notable categories included post-translational modification, protein-turnover, and chaperones (n = 97); signal-transduction mechanisms (n = 96); nucleotide-transport and metabolism (n = 86); and defense mechanisms (n = 70). Additionally, genes associated with Mobilome, prophages, and transposons (n = 66); cell-cycle control, cell division, and chromosome maintenance (n = 43); secondary metabolites biosynthesis, transport, and catabolism (n = 33); intracellular-trafficking, secretion, and vesicular-transport (n = 26); cell-motility (n = 7); extracellular-structure and cytoskeleton (n = 3); and other categories (n = 1) were identified. These results indicate that strain d1418m22 may have resistance to stress conditions.

**Figure 3.**
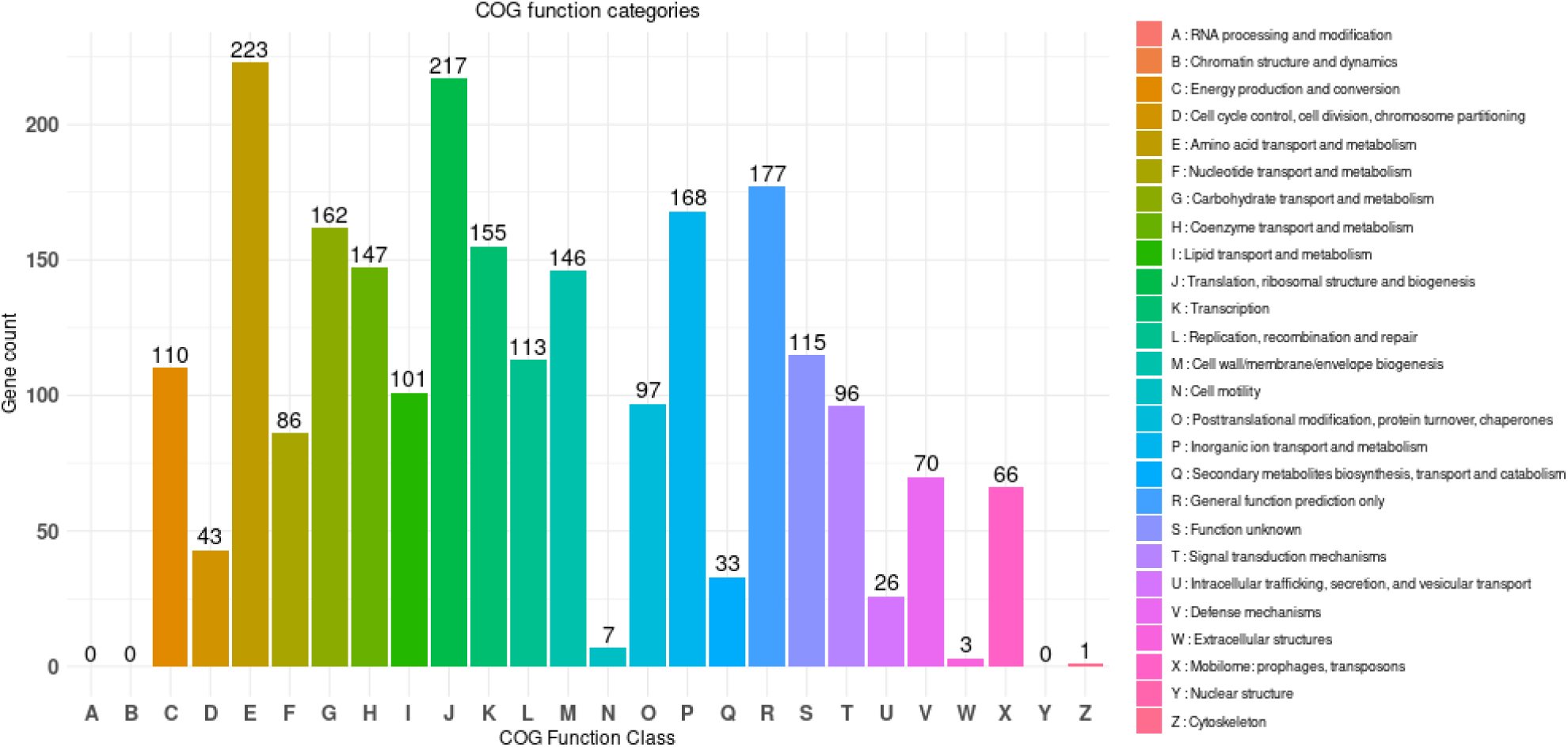
RAST subsystems categories and feature distribution of MRSA strain d1418m22 genome.

A similar study investigated 60 MRSA isolates, with 40 obtained from patients and 20 from food samples. Analysis based on homologous whole-cluster COG expansions revealed that 1,040 genes were involved in metabolism, including amino-acid-transport, carbohydrate-transport, and inorganic ion transport. Additionally, 564 genes were linked to information storage and processing, which encompassed processes like transcription, replication, recombination, repair, translation, and ribosomal structure. Furthermore, 396 genes were associated with cellular processes and signalling, including cell wall, membrane-envelope, biogenesis, protein-turnover, chaperones, post-translational modifications, and defence mechanisms (Liao et al., 2020).

### 3.5. Gene Ontology (GO) and KEGG pathway enrichment analyses were performed to explore the functions and biological pathways of the identified genes.

Gene ontology (GO) annotation of proteins from the non-redundant (NR) database was carried out using OmicsBox to classify genes into three domains: biological processes, molecular functions, and cellular components. In the MRSA strain d1418m22 genome, 801 genes were assigned GO terms (Supplementary Table S1, Figure 4). Prodigal predicted 2,625 protein-coding genes, of which 1,961 were confirmed as homologous to *S. aureus* using BLASTP with an E-value threshold of 1e^−6^. The GO terms included 1,286 for biological processes, 423 for cellular components, and 1,466 for molecular functions. A BLASTP homology search (E-value ≤ 1e^−5^) further identified the 20 antibiotic resistance genes by comparing the *S. aureus* genome to the Comprehensive Antibiotic Resistance Database (CARD).

**Figure 4.**
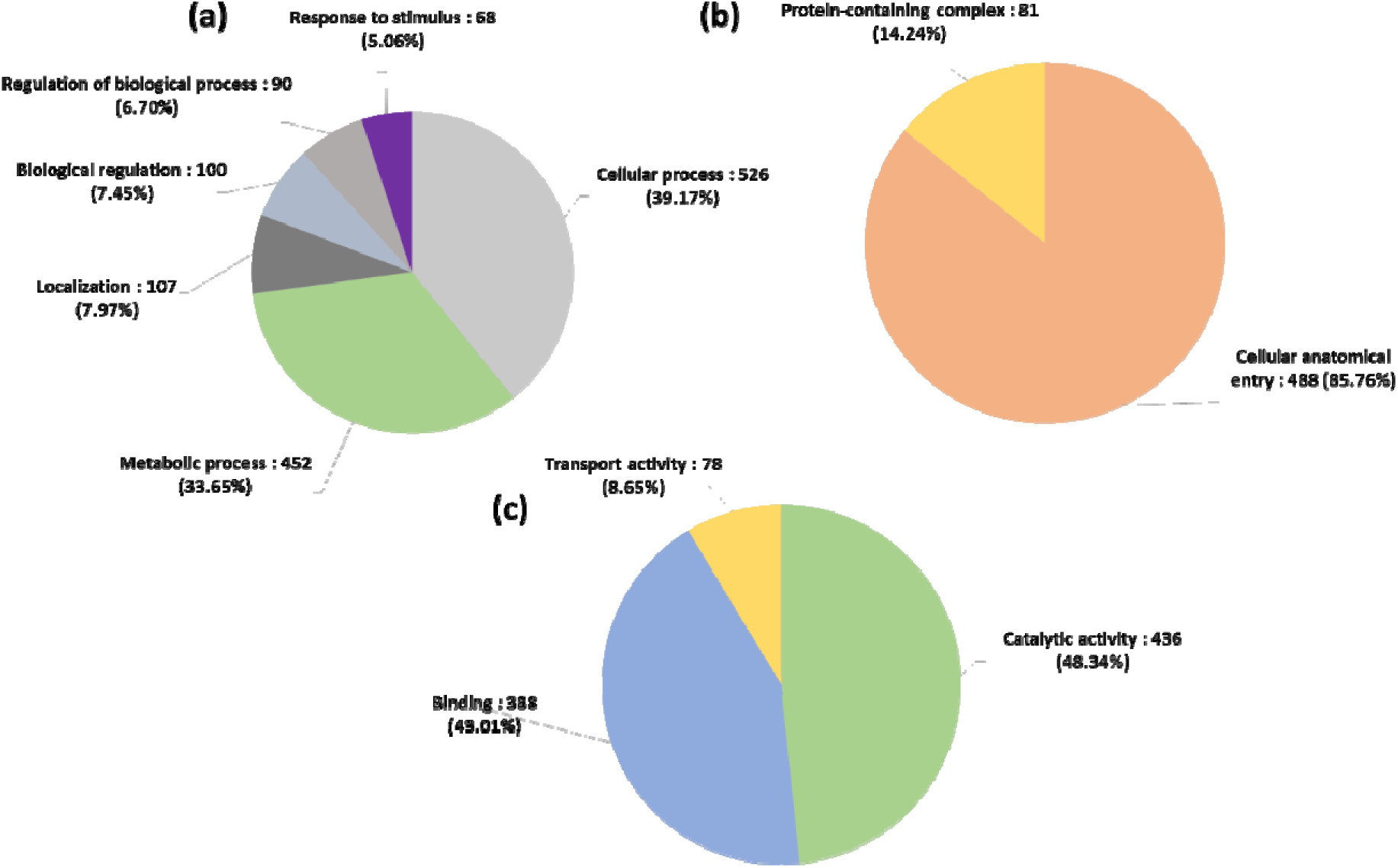
**(a)** GO (Biological Process) distribution for **strain d1418m22 (b)** Cellular Component distribution **(c)** Molecular Function.

Pathway analysis, ortholog-assignment, and gene mapping were conducted using the KEGG Automatic Annotation Server (KAAS), as summarized in Table 3. Of the 2,625 predicted gene sequences from the strain d1418m22 genome, 1,590 were mapped to KEGG pathways via BLASTP (bit-score ≥ 60), covering metabolic pathways related to carbohydrates, lipids, nucleotides, amino acids, glycans, cofactors, vitamins, and others. Key pathways included carbohydrate metabolism (296 proteins), amino acid metabolism (199), metabolism of cofactors and vitamins (127), energy metabolism (107), membrane transport (117), signal transduction (93), cellular community—prokaryotes (64), replication and repair (74), and translation (82). These pathways involved genes associated with genetic and environmental information processing as well as cellular processes. In a related study, approximately 65% of mutants causing endothelial cell (EC) damage were linked to metabolism, 24% to environmental information processing, 11% to genetic information processing, and 9% to cellular processes, with some genes displaying multifunctionality (Xiao et al., 2022).

### 3.6. Identified antibiotics resistant, virulence genes and CRISPRC as from MRSA d1418m22 genome

Advancing computational tools and databases for *in silico* AMR prediction from WGS data are crucial for AMR surveillance. Using the CARD, 20 AMR genes were identified in the assembled d1418m22 genome. Details on resistance mechanisms and gene functions are provided in Supplementary File S1.

Our integrated approach, combining bioinformatics tools, provided a detailed analysis of AMR and virulence determinants, with stricter identity thresholds than RESFinder and CARD, ensured precise detection of resistance and virulence features. The CARD-RGI tool effectively identified resistance genes and their identity scores in the assembled genomes. Detected genes included *mgrA* (100%), *mepR* (100%), *kdpD* (99.89%), *arlR* (100%), *arlS* (100%), *tet(38)* (99.78%), *norC* (99.57%), *mecR1* (100%), *mecA* (99.7%), *dfrC* (98.14%), *AAC(6’)-Ie-APH(2’’)-Ia* biofunctional protein (100%), *S. aureus parC* (99.5%), and *S. aureus gyrA* (99.66%), conferring resistance to fluoroquinolones. These findings are presented in Figure 5 (a), with additional details in Supplementary File S1. CARD-RGI provided more comprehensive results, identifying multiple efflux pumps, antibiotic inactivation, target replacement, and alteration proteins linked to antibiotic resistance, as detailed in Supplementary File S1. Kmiha, (2023) reported high resistance to β-lactam, quinolone, and aminoglycoside antibiotics, with cefoxitin-(100%), ciprofloxacin-(86%), gentamicin-(83%), and penicillin-(75%) showing the highest levels. Moderate to low resistance was observed for Fosfomycin-(28%), oxacillin-(26%), fusidic-acid (25%), amikacin-(23%), tobramycin-(22%), ampicillin-(22%), kanamycin-(20%), tetracycline-(17%), erythromycin (8%), and teicoplanin-(1.5%).

**Figure 5.**
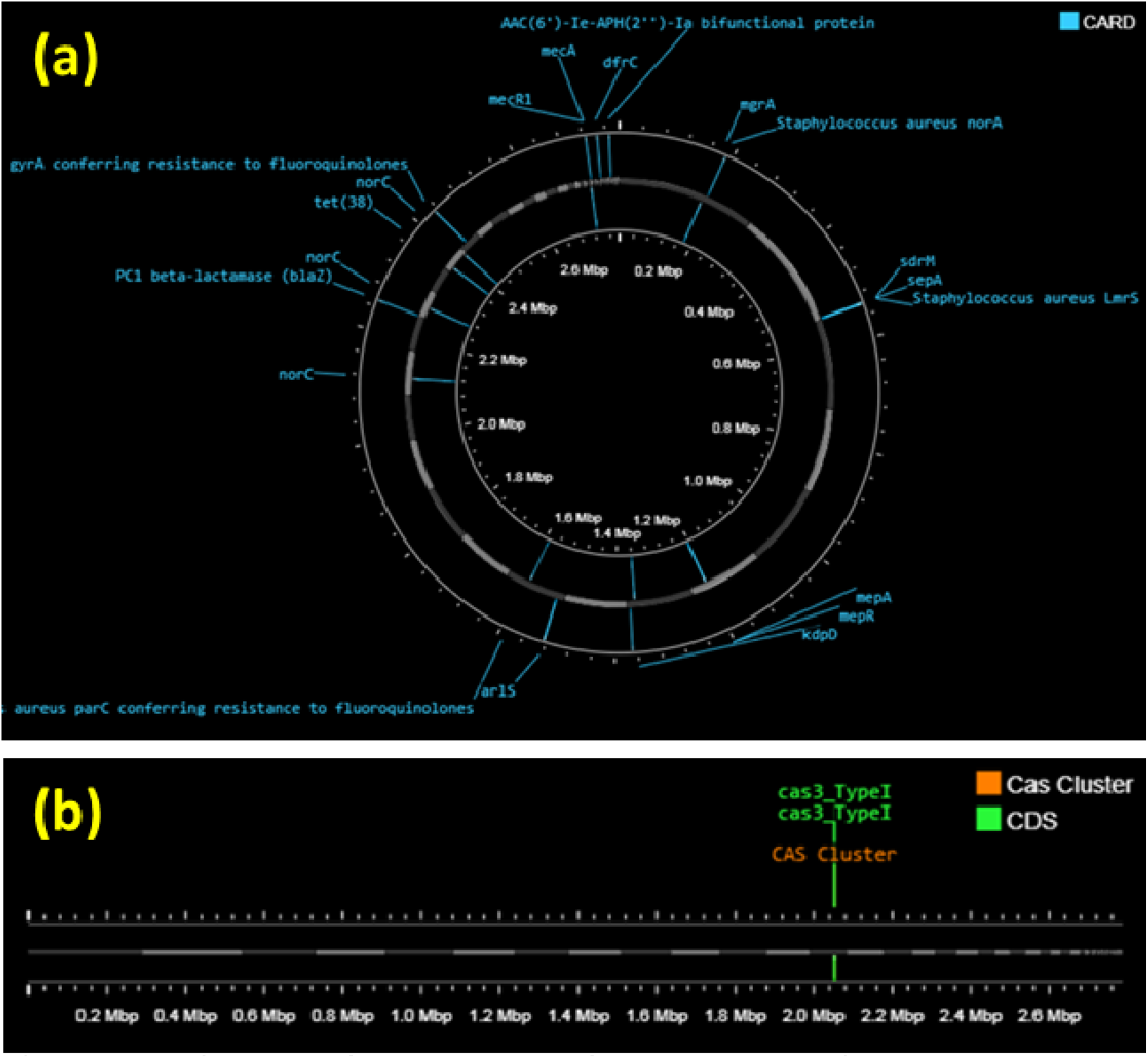
**(a)** The CARD **(b)** CRISPRCas Finder in MRSA strain d1418m22 strain genome (Prokka 1.14.6 tool).

Thus, besides being an MDR strain, MRSA d1418m22 also harbors numerous virulence genes, a finding that is similar to many other MRSA strains that have been reported to carry various virulence and resistance determinants that are crucial for the success of this bacterium in the human host (Sharma et al., 2017; McClure et al., 2018). Further in our investigation, resistant MRSA d1418m22 strain identified key resistance genes included mgrA, mepR, arlR, arlS, mecR1, and AAC(6’)-Ie-APH(2’’)-Ia (all 100% identity), along with others like mecA (99.7%), norC (99.57%), tet(38) (99.78%), and blaZ (94.66%). In six reference strains, eye-isolated <u>GCA_015219905.1</u> carried dfrG, AAC (6’)-Ie-APH (2’’)-Ia, arlR, and sdrM, while <u>GCA_015219885.1</u>), exhibited similar genes. In case of wound pus isolated resistant strains, wound/pus-isolated <u>GCA_033738955.1</u> included dfrG, mepA, norC, mecI, and APH(3’)-IIIa, whereas <u>GCA_000695215.1</u> carried mecR1, mepR, and mgrA. Additionally, urine-isolated strains, <u>GCA_022693325.1</u> and <u>GCA_022693305.1</u>, shared resistance genes like mecR1, tet(38), mepR, vanA cluster genes (vanR, vanH, vanZ), and AAC(6’)-Ie-APH(2’’)-Ia, highlighting significant resistance profiles. Seven genes encoding efflux pumps from the major-facilitator-superfamily-(MFS), small-multidrug-resistance-(SMR) pumps, and multidrug and toxic compound extrusion (MATE) transporters were identified in the d1418m22 genome. These genes, including *mgrA* (100%), *mepR* (100%), *kdpD* (99.89%), *arlR* (100%), *arlS* (100%), *tet (38)* (99.78%), and *norC* (99.57%), are associated with resistance to fluoroquinolones, glycylcyclines, and tetracycline antibiotics. Previous studies (Ikonomidis et al., 2009; Holtfreter et al., 2007) have highlighted the role of these efflux-pumps in enabling resistance to antiseptics and disinfectants antibiotics. Additional, in our result resistance determinants identified on the d1418m22 genome chromosome include *mecR1* (100%), *mecA* (99.7%), *dfrC* (98.14%), and the *AAC-(6’)-Ie-APH-(2’’)-Ia* biofunctional protein (100%), conferring resistance to penam, diaminopyrimidine, and aminoglycoside antibiotics. Pillai, (2002) revealed that, the tetracycline resistance determinant *tet-(38)* resistance gene was consistently detected in all genomes, while *tet-(K)* and *tet-(M)* were less frequently identified. The observed limited tetracycline resistance suggests it may be associated with mobile *tet*-genes, such as *tet-(K)* and *tet-(M)* (Pillai et al., 2002). Similarly, ciprofloxacin resistance arises from mutations in *gyrA*, *grlA*, or *grlB* (Schmitz et al., 1998). Furthermore, *Staphylococcus aureus parC* (99.5%) and *Staphylococcus aureus gyrA* (99.66%) were detected, both associated with fluoroquinolone resistance. Lakhundi, (2018) found that 78 percent of MRSA isolates carried the *mecA*-gene, while 22 percent were *mecA*-negative, indicating the potential presence of alternative cassette genes such as *mec-C*, *mec-B*, or *mec-D*. The spread of *mecA*-positive MRSA in clinical sites has been observed in many African and European countries, with the resistance being transmitted via *SCCmec* cassette genes through HGT (Baig et al., 2018).

In our result, in addition to chloramphenicol and erythromycin, most phenotypic resistance and acquired AMR genes were identified using the ResFinder-4.6.0 server. The resistance genes *aac-(6’)-aph-(2’’)*, *mecA*, and *blaZ* were found to confer resistance to aminoglycosides, quinolones, and beta-lactams. The acquired AMR gene hits and their identity scores were *aac(6’)-aph(2’’)* (100%), *blaZ* (99.98%), and *mecA* (100%), as detailed in Supplementary File S1.

Previous studies have also shown that antimicrobial inactivation by enzymes, such as the modification of aminoglycosides by aminoglycoside-modifying-enzymes (AMEs) encoded by genes like *aac-(6’)/aph-(2’’)*, *aph-(3’)-IIIa*, and *ant-(4’)-Ia*, contributes to resistance (Khosravi et al., 2017). Similarly, the bio-functional AME gene *[aac(6)-Ie-aph(2)-Ia]* was noted in all MRSA isolates. This gene imparts significant resistance to aminoglycosides like gentamicin, but does not affect streptomycin (Livermore, 2000). The *aac-(6)-aph-(2)*-gene, which encodes an aminoglycoside-acetyltransferase conferring gentamicin-resistance, is located on the *SCCmec* element. Additionally, the lmrS-gene, encoding MFS-transporter, might contribute to *SauR3*’s decreased susceptibility to macrolides and aminoglycosides (Khosravi et al., 2017).

### Virulence factors

Twenty-one VFs were detected in the d*1418m22* strain genome, along with their identity scores: *sak* (100%), *scn* (100%), *aur* (99.93%), *hlgA* (100%), *hlgB* (100%), *hlgC* (100%), *lukF-PV* (100%), *lukS-PV* (100%), *sec* (99.88%), *seg* (99.87%), *sei* (99.73%), *sel* (99.72%), *sem* (99.86%), *sen* (99.74%), *seo* (100%), *seu* (100%), and *tst* (100%). The VFs determinants were classified into the following categories: Staphylokinase, Staphylococcal complement inhibitor, Aureolysin, Gamma-hemolysin (encompassing the chain II precursor, component B precursor, and component C), PVL (including F and S components, with duplicates), and enterotoxins (C, G, I, L, M, N, O, U), in addition to toxic shock syndrome toxin-1. Detailed information is provided in Supplementary File S1.

The genome of the d1418m22 strain was confirmed to possess the PVL gene (lukS/F-PV), located on an SA2PM1-like phage that exhibited 100% identity with the SAPM1 phage from the *S. aureus* PM1 strain (Supplementary File S1). Furthermore, the d1418m22 strain genome revealed 21 VFs with 100% identity, including sak, scn, hlgA, hlgB, hlgC, lukF-PV, lukS-PV, seo, seu, and tst.

In six resistant reference strains, VFs varied based on the source of isolation. For eye-isolated strains (<u>GCA_015219905.1</u> and <u>GCA_015219885.1</u>), the identified VFs included scn, hlgA, hlgB, hlgC, lukF-PV, lukS-PV, sea, sec, sec3, sen, seo, seb, and sak, all showing 100% identity. In wound pus-isolated strains (<u>GCA_033738955.1</u> and <u>GCA_000695215.1</u>), the detected VFs included sak, scn, aur, hlgA, hlgB, hlgC, sen, seo, and seu. For urine-isolated strains (<u>GCA_022693325.1</u> and <u>GCA_022693305.1</u>), shared VFs comprised hlgA, hlgB, hlgC, lukD, lukE, sem, seo, aur, splA, and splB, all with 100% identity. Our findings indicate that MRSA strains carry a wide range of VFs, with enterotoxin genes being particularly prevalent, suggesting their spread among strains with similar genetic backgrounds (Xu et al., 2021). The lukF/S-PV gene was detected in 6.9% of *S. aureus* isolates from diabetic foot infections in a study by Viquez-Molina, (2018), while a study in Lisbon did not find this gene in the same context (Mottola et al., 2016). The PVL locus, which resides on a bacteriophage, is linked to skin infections and, in some instances, severe necrotizing pneumonia (Lina et al., 1999; Gillet et al., 2002).

The PVL gene in MRSA strains is mainly associated with skin and soft tissue infections, especially in community-associated clones (Abdulgader et al., 2015). The d1418m22 strain genome revealed the presence of the cas3 Type I cluster (Figure 5b), spanning 1347 bp on contig Scaffold_13. The cas3 gene is a crucial part of the Type I CRISPR-Cas system, which allows *S. aureus* to recognize and neutralize foreign genetic elements, such as plasmids and bacteriophages, thereby ensuring genomic stability. The SCCmec is a MGE contains the mecA gene, responsible for resistance to β-lactam antibiotics. The mecA, mecR1, and subtype-IVa (2B) genes were identified in the d1418m22 strain with identity percentages of 100%, 100%, and 99.93%, respectively. These findings suggest the potential existence of additional, unidentified regulators of the mecA gene, which could lead to a revision of the mecA regulatory mechanism in this strain. Further details are provided in Supplementary File S1. In a comparable study, the MRSA phenotype was linked to the mecA gene, which encodes penicillin-binding protein (PBP2a), a protein with reduced affinity for β-lactams. The mecA gene is located within the SCCmec element, and some MRSA strains also contain regulatory genes such as mecI and mecR1, which modulate mecA expression (Oliveira et al., 2011). At Charles Nicole Hospital in Tunis, SCCmec types I, II, and III were predominantly observed (Jemili-Ben et al., 2006), whereas SCCmec type IV was most frequently found in clinical studies across Tunisia (Abdulgader et al., 2015; Ben et al., 2013; Kechrid et al., 2011; Bouchami et al., 2009).

### 3.7. Genomic features of MRSA d1418m22 genome

A total of 2,625 genes were identified in the d1418m22 genome, comprising 2,583 coding sequences (CDSs), 56 tRNAs and 6 rRNAs. The genomic distribution of protein-coding sequences, rRNA, tRNA genes, and GC skew is depicted in Figure 6. The genome of MRSA strain SO-1977 has been previously described as spanning 2,827,644 base pairs (bp) with a GC content of 32.8%, and comprising 59 RNAs and 2,629 predicted CDSs (Ali et al., 2019). Additionally, a study on *S. aureus* ST22, particularly the EMRSA-15 epidemic strain, reported a genome size of 2,831,239 bp, with a GC content of 32.7%, and a total of 2,835 genes, including 2,768 CDSs, 56 tRNAs, and 7 rRNAs (Ullah et al., 2022). The genome of d1418m22 was assembled into 61 scaffolds, with an N50 value of 130,908 bp, the largest scaffold measuring 290,662 bp. In contrast, Ullah (2022) reported that the *de novo* assembly of the ST22 genome resulted in 52 contigs longer than 500 bp, with the largest contig measuring 425,597 bp. In our study, a total of 2,583 CDSs were identified, with 870 encoding “hypothetical proteins” of unknown function. The remaining genes are primarily associated with carbohydrate metabolism and stress response pathways. These genes are notably linked to various key proteins, such as the HTH-type transcriptional regulator GabR, protein-arginine kinase activator, DNA repair protein RadA, transcriptional regulator NusG, bacitracin export ATP-binding protein BceA, cytokinin riboside phosphoribohydrolase, fatty acid resistance protein FarB, amino-acid ABC transporter-binding protein, L-cystine permease TcyB, NADP-dependent oxidoreductase YfmJ, zinc-type alcohol dehydrogenase-like protein, signal recognition particle protein, L-cystine import ATP-binding protein TcyC, KDP operon regulatory protein KdpE, conserved virulence factor B, oligoendopeptidase F, and plasmid (Figure 6 & Supplementary File S1).

**Figure 6.**
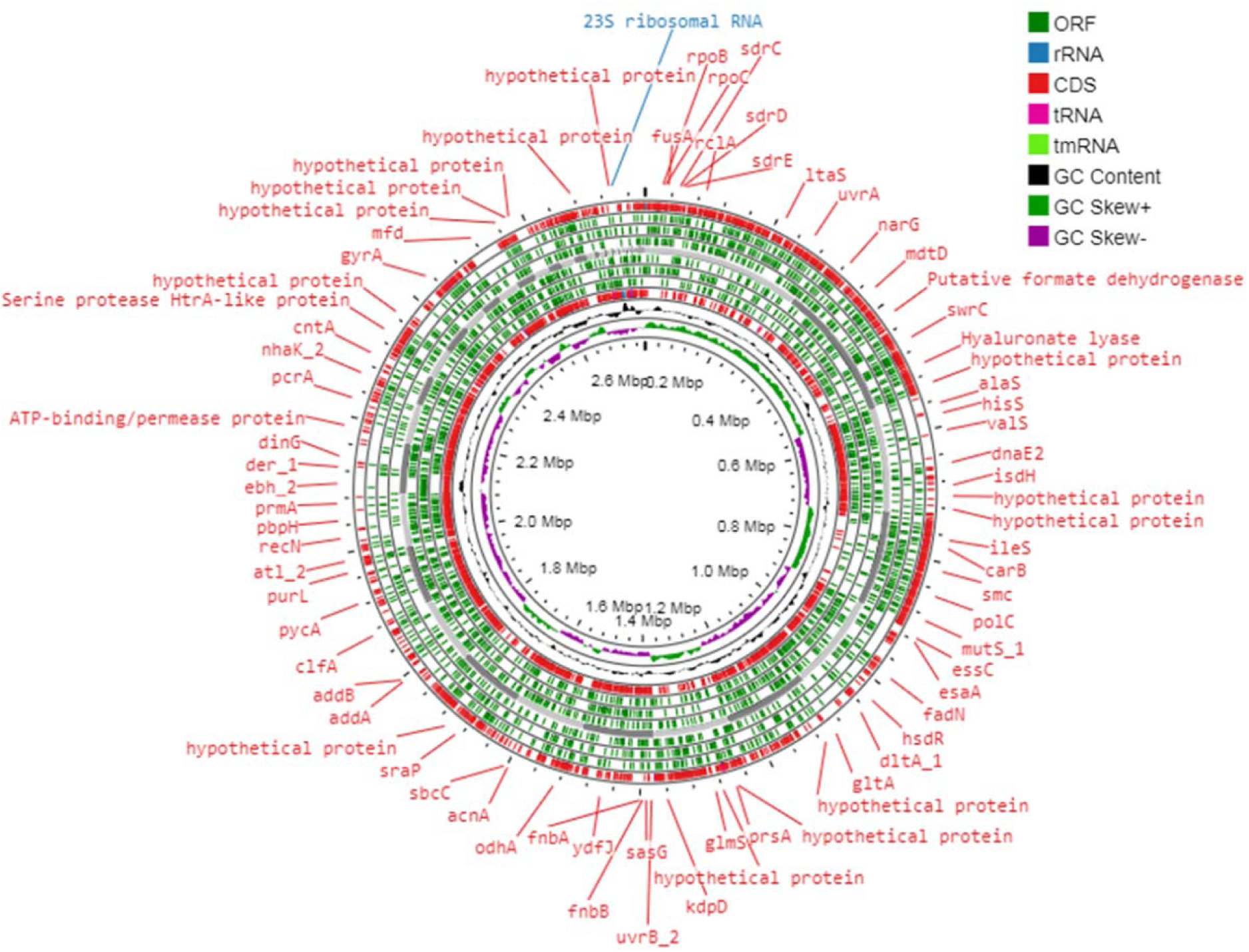
The circular map of the MRSA d1418m22 genome shows CDSs in the outer red ring, with ORFs, tRNA, and rRNA genes in the inner green ring, representing the positive and negative strands in circles 1 and 2, respectively. Circle 3 (black and green) depicts the GC skew, while circle 4 (purple) represents the deviation in GC content from the genome’s average.

### 3.8. Mobile genetic elements in MRSA d1418m22 genome

MGEs in MRSA refers to genome segments acquired from other organisms through transformation, transduction, or conjugation. These regions typically harbor genes that confer advantages, including antibiotic resistance or increased virulence (Lindsay et al., 1998). In this study, 106 HGT regions were identified in the d1418m22 strain, as shown in Figure 7. These genes demonstrate the role of HGT in enhancing the strain’s metabolic adaptability and resilience to environmental pressures. A detailed analysis uncovered 237 MGEs within the d1418m22 genome, categorized as follows: integration/excision (64), replication/recombination/repair (80), phage-related (48), stability/transfer/defense (18), and transfer (27) (Figure 7 and Supplementary File S1). Annotation using mobileOG-db with an E-value threshold of 1.0e-05 revealed major categories such as phage-related functions, replication/recombination/repair, transfer, integration/excision, and stability/transfer/defense. Minor categories included replication, infection, competence, conjugation, infection regulation, initiation, copy number control, plasmid replication, chaperone functions, partitioning, and lysis/lysogeny (details in Figure 7 & Supplementary File S1). A comparative analysis with a previous study on MRSA CC22 identified 37 distinct MGEs, including SCC elements, pathogenicity islands, transposons, and plasmids. Key findings from the prior study included the widespread presence of a Tn552-like transposon carrying the *blaZ* gene (penicillin resistance) and SCCmec IV (methicillin resistance). The SaPIsec pathogenicity island, encoding the *sec* gene for enterotoxin C, was found in 61% of isolates and exhibited high similarity to a region in the reference genome. Additionally, two plasmids were identified: P1-ermC, a 2.5 kb plasmid carrying the *ermC* gene for erythromycin resistance, and P2-hm, 32 kb plasmid harboring genes for heavy metal resistance (Jamrozy et al., 2017). Additionally, six reference strains were identified with SCCmec elements. In the eye-isolated resistant strains (<u>GCA_015219905.1</u> and <u>GCA_015219885.1)</u>, the predicted whole cassettes and % template coverage was SCCmec type V (5C2) at 50.96% and SCCmec type IVc (2B) at 78.26%, respectively. For wound pus-isolated strains (<u>GCA_033738955.1</u> and <u>GCA_000695215.1)</u>, the predicted whole cassettes and % template coverage was SCCmec type III (3A) at 94.82% and SCCmec type IV (2B) at 97.25%. Similarly, in urine-isolated resistant strains (<u>GCA_022693325.1</u> and <u>GCA_022693305.1),</u> the predicted whole cassettes and % template coverage was SCCmec type IV (2B) at 79.86% and 79.79%, respectively. In the resistant d1418m22 strain genome, the predicted whole cassette and % template coverage was identified as SCCmec type IVa (2B) at 94.66%.

**Figure 7.**
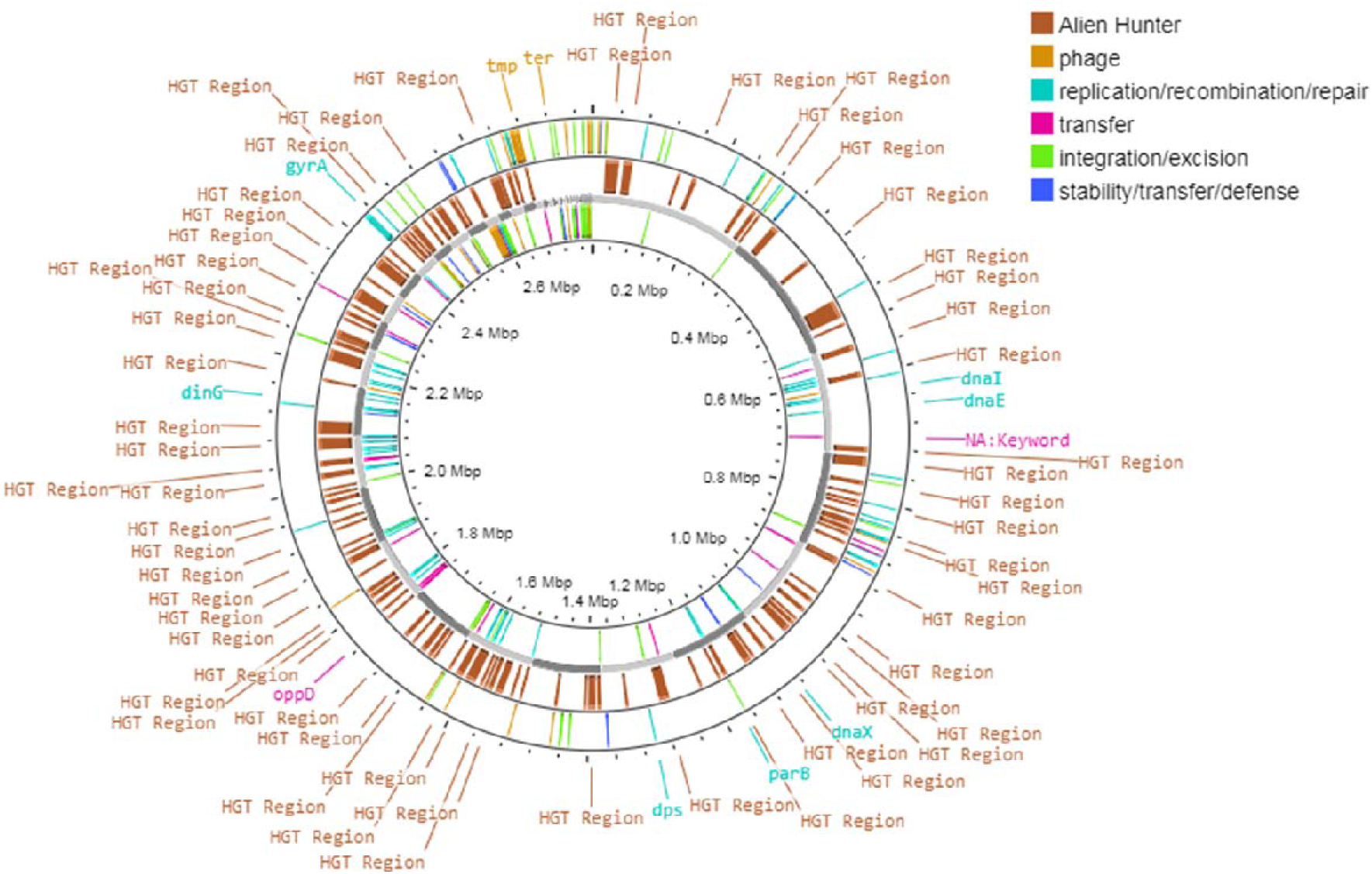
Identify open reading frames (ORFs), rRNA, CDS, tRNA, and tmRNA, and annotate the genome sequence using the Prokka tool (version 1.14.6).

### 3.9. Characteristic of prophages-like elements

Prophage regions in the d1418m22 strain are segments of the genome where bacteriophage DNA has integrated. These prophages contribute to the bacterium’s pathogenic traits, such as virulence, antibiotic resistance, genetic diversity, biofilm formation, and immune evasion. Phage screening using PHASTER revealed two prophage insertions in the d1418m22 genome, both intact but categorized as incomplete prophages (Figure 8). These intact prophages, located at Scaffold_23 and Scaffold_26, are 25.5 kb and 23.9 kb in size, respectively, containing 84 CDSs. Of these, 22 are hypothetical proteins of unknown function, while 29 are common phage-related proteins, including endonuclease, head, capsid, tail, integrase, portal protein, terminase, and protease, as shown in Figure 8. A related study by Naorem et al. (2021) identified three prophage regions in *S. aureus* strain SA G6, including an intact prophage (phiG6.2, 72.8 kb), a questionable prophage (phiG6.3, 74.5 kb), and an incomplete prophage (phiG6.1, 16.4 kb).

**Figure 8.**
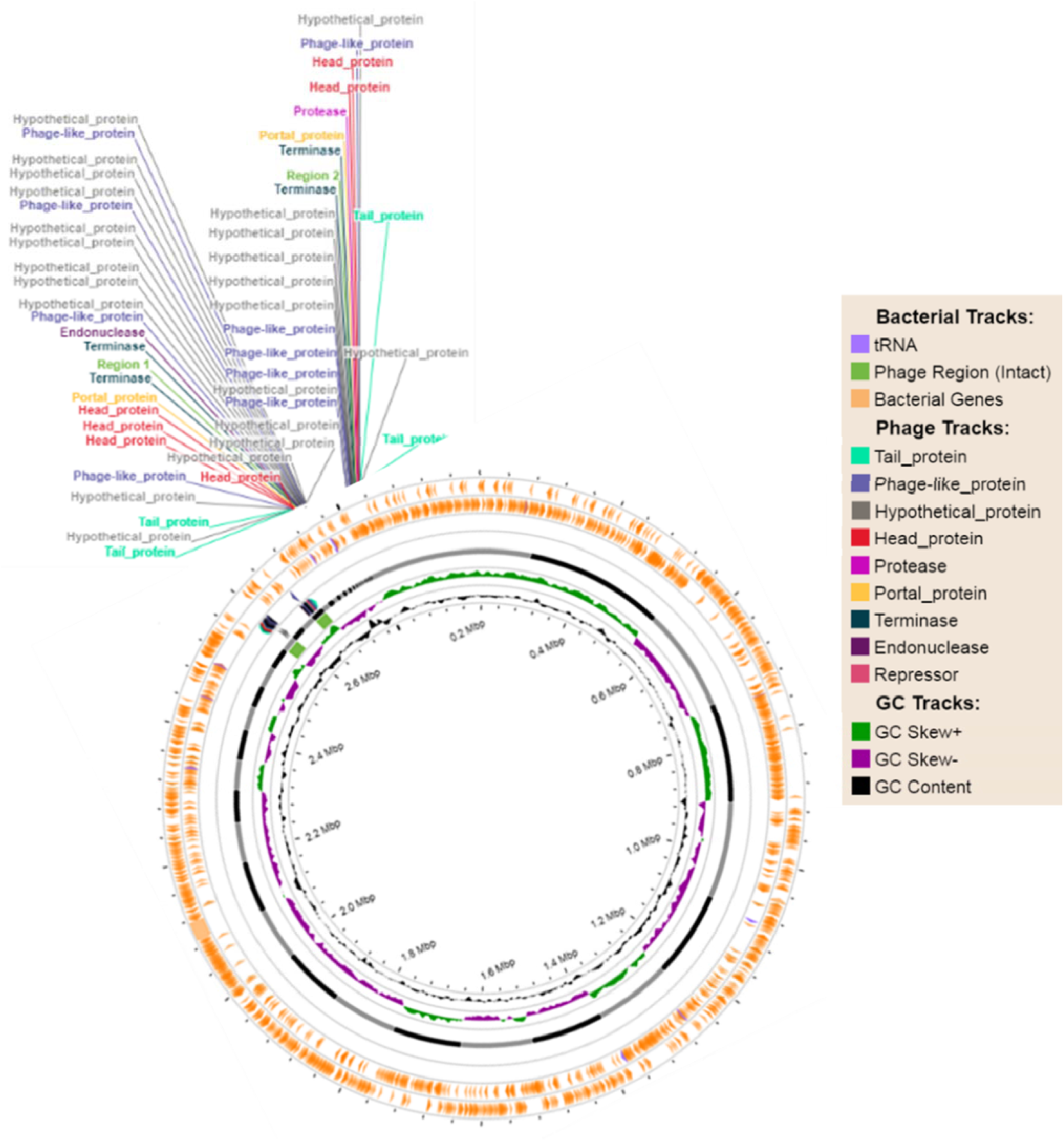
MRSA strain d1418m22, Prophage regions have been identified, of which 2 intact prophage regions are (25.5 and 23.9 kb).

### 3.10. BLAST comparison with references strains and ANI

Perform a BLAST comparison with another genome sequence by *BLAST+2.15.0 tool.* The six reference MRSA strains, MRSA-AMRF4 (<u>GCA_015219905.1</u> and MRSA-AMRF 5 (<u>GCA_015219885.1)</u>, were isolated from eye infections, both with a genome size of 2.9 Mb and 33% GC content. MRSA (<u>GCA_033738955.1)</u> and MRSA-15 (<u>GCA_000695215.1)</u>, isolated from wound pus samples, have genome sizes of 3 Mb and 2.8 Mb, respectively, with 33% GC content. Additionally, VMRSA-WC071 (<u>GCA_022693325.1</u>) and VMRSA-WC081 (<u>GCA_022693305.1</u>), isolated from urine, also have 2.9 Mb genomes and 33% GC content. In our resistant MRSA strain d1418m22 shares similar characteristics, with a 2.78 Mb genome and 32.8% GC content showed in figure 9. The FastANI 1.34 tool was used for fast, alignment-free computation of whole-genome Average Nucleotide Identity (ANI). The reference strains *MRSA* <u>GCA_015219905.1</u> and *MRSA* <u>GCA_015219885.1</u> exhibited ANI values of 98.19 and 98.11, respectively, when compared to the resistant *MRSA d1418m22* genome. For wound pus-isolated genomes, the ANI values were 98.12 and 99.80 for the bacterial strains <u>GCA_033738955.1</u> and *MRSA* <u>GCA_000695215.1</u>, respectively, in comparison with the *d1418m22* genome. Additionally, urine-isolated bacterial strains (*MRSA* <u>GCA_022693325.1</u> and *MRSA* <u>GCA_022693305.1</u>) demonstrated ANI values of 98.12 and 98.16, respectively, when compared to the *d1418m22* genome (as shown in Figure 10).

**Figure 9.**
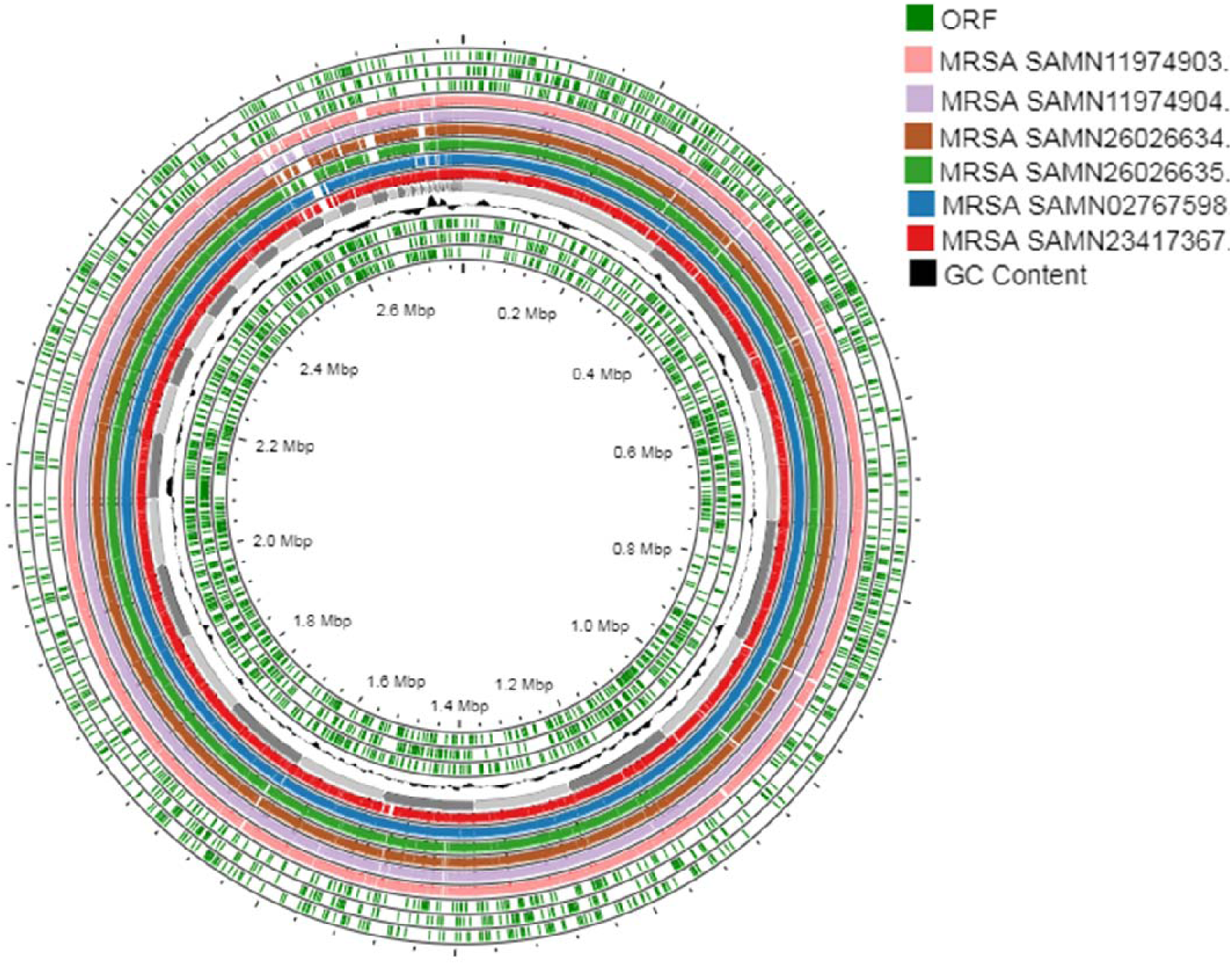
Comparison of the *S. aureus* d1418m22 genome sequence using CGView Proksee with genomes of closely related MRSA d1418m22 strain, i.e., MRSA-AMRF4 (<u>GCA_015219905.1</u>) and MRSA-AMRF5 (<u>GCA_015219885.1</u>), MRSA (<u>GCA_033738955.1</u>) and H-EMRSA-15 (<u>GCA_000695215.1</u>), VMRSA-WC071 (<u>GCA_022693325.1</u>) and VMRSA-WC081 (<u>GCA_022693305.1</u>) strains. The MRSA d1418m22 genome was used as the main reference genome for the CGView Proksee BLASTN comparison. From outside to the center, rings 1 and 3 show (green) ORF (Open Reading Frame) of three strain and in side 3 rings (Green) also ORF. Inside light and dark gray color ring showed coding sequences for the MRSA d1418m22 forward and reverse strands; from outside ring 4 shows the GCA_015219885.1 genome (in pink), ring 5 shows GCA_015219905.1 (in purple), ring 6 shows GCA_022693325.1 (in sky brown), ring 7 shows GCA_022693305.1 (in green), ring 8 shows GCA_000695215.1 (in blue), ring 9 shows GCA_033738955.1 (in red) and finally, the innermost ring 10 shows a plot of the GC content.

**Figure 10.**
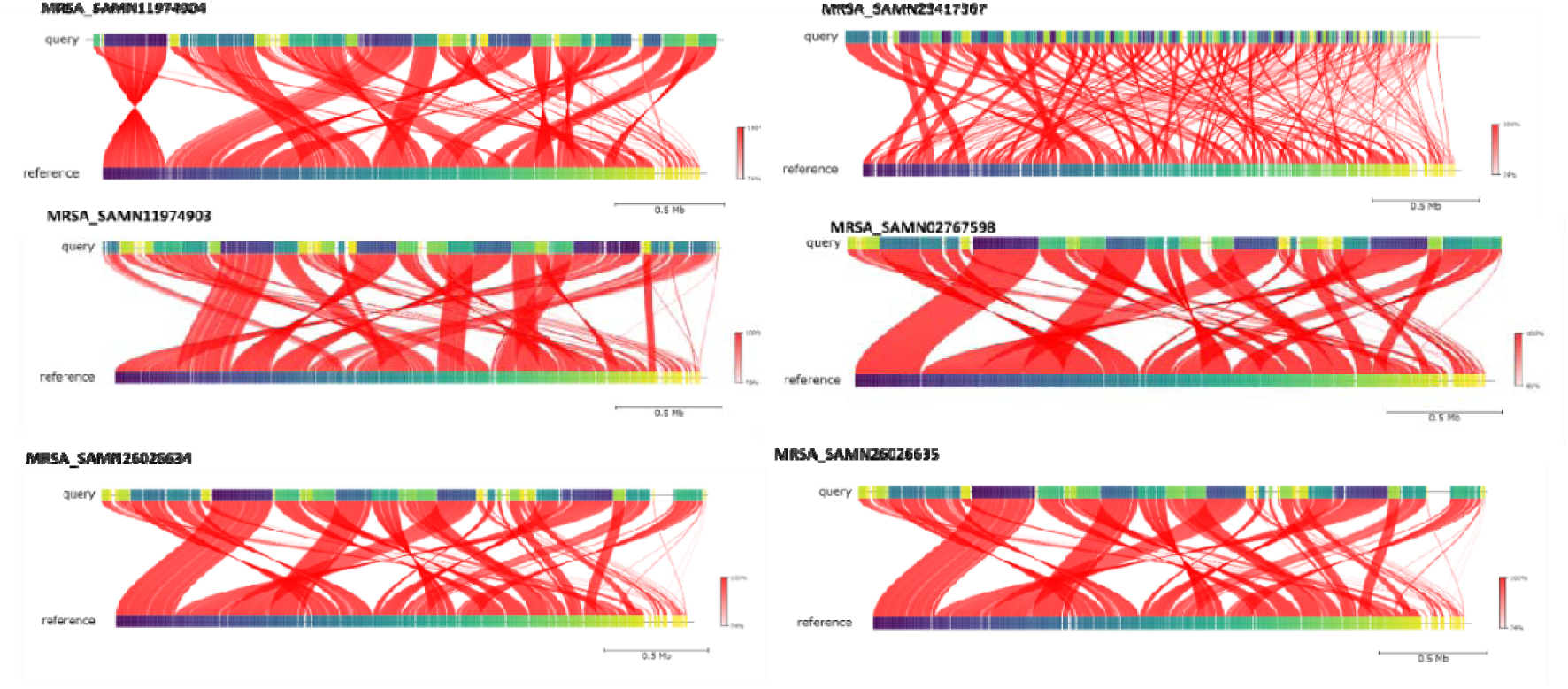
The locations of the genomic average nucleotide Identity (ANI) identified in the MRSA d1418m22 strain with references genome are labelled. MRSA-AMRF4 (<u>GCA_015219905.1</u>) and MRSA-AMRF5 (<u>GCA_015219885.1</u>), MRSA (<u>GCA_033738955.1</u>) and H-EMRSA-15 (<u>GCA_000695215.1</u>), VMRSA-WC071 (<u>GCA_022693325.1</u>) and VMRSA-WC081 (<u>GCA_022693305.1</u>) strains.

### 3.11. Interaction Network Analysis of hub genes

The 2,625 predicted genes from the MRASA strain d1418m22 genome were analyzed for similarity using the BLASTP algorithm against NCBI’s non-redundant (nr) database, applying an e-value cutoff of 1e^-5^. Out of 2625 unique genes, 1835 genes showed the highest interaction and neighborhood score identified in d1418m22 strain. The neighborhood score 1835 genes of identified hub genes ranges 0.99 detailed information in Supplementary S2. The STRING score is useful for identifying hub genes in the MRSA d1418m22 strain, which are genes that have a high degree of interaction with other genes in a network. Figure (11 a) showed the network of all 2625 genes, these genes serve as hubs in the network. In this network the STRING network, the 1835 identifies hub genes because they have thicker or more numerous connections in the network (Fig. 10a). Out of 1835 highest score hub genes, eight genes identified as biofilm formation in MRSA d1418m22 strain and showed high interaction to each other such as *eno- pgk* (0.99), *eno- pyk* (0.99), *eno*- *pckA* (0.96), *eno-fda* (0.94), *eno-zwf* (0.92), *eno-rplD* (0.91), *eno-pfkA* (0.90), *icaA-icaB* (0.99), *icaA-icaD* (0.97), *icaA-icaC* (0.97), *icaC-icaD* (0.96), *icaB-icaD* (0.96), *icaB-icaC* (0.96), *icaA-icaR* (0.93), *icaD-icaR* (0.93), *fnbA- spa* (0.96), *clfA- fnbA* (0.90) *eno- tpiA* (0.99), *eno- gpmI* (0.99), *eno- pgi* (0.99) showed list of genes in Supplementary file S2. The regulatory network leading biofilm formation and virulence in MRSA involves complex interactions among various systems that contribute to its pathogenicity (Guo et al., 2020). A similar study by Atshan, (2013) identified 13 biofilm-related genes using PCR with specific primers and established conditions. Spa-typing is a highly effective tool for epidemiological studies of Staphylococcus aureus, offering excellent discriminatory power. It targets sequence variations in the hypervariable region of the spa gene, which encodes staphylococcal protein A, essential for adhesion and colonization. The prevalence of spa types varies across regions and sources (Abdulgader et al., 2015; Hara et al., 2016). The identified genes included icaA, icaB, icaC, and icaD (involved in intercellular adhesion); fnbA and fnbB (associated with fibronectin binding); clfA and clfB (clumping factors); cna (collagen-binding protein); eno (laminin-binding protein); ebpS (elastin-binding protein); fib (fibrinogen-binding protein); and bbp (bone sialoprotein-binding protein) (Atshan et al., 2013). Notably, icaA, icaD, fnbA, and eno were most commonly identified, with icaA and icaD playing essential roles in intercellular adhesion and biofilm development in *S. aureus* (Nourbakhsh et al., 2016). According to Cerca et al., (2018), it also interacts with *icaR*, a gene whose protein inhibits biofilm formation. The icaADBC operon is regulated by global factors, including SarA and σB, which exert indirect influence, as well as direct regulation by IcaR (Cue et al., 2012).

We utilized tools within the Cystoscape software to identify network hubs, which have the highest number of interactions (edges) within the network. Both cytoHubba and MCODE produced identical results in identifying these hub genes. Figure 11 b, c, and d displays the top-scoring clusters containing 10, 30, and 50 nodes, respectively. A detailed list of the top 10, 30, and 50 hub genes is provided in Supplementary File S2. In this network top 10 genes identified as high degree interaction by other genes such as guaA (interaction with 232 genes), rpoB (182), metG (178), pheT (176), hisF (169), rplB (154), rpoA (152), rplM (152), infB (150) and fusA (150). The interaction network of top 13, 30 and 50 genes showed in Figure 11 b, c & d. These genes are involved in essential cellular functions like transcription, translation, and nucleotide/amino acid metabolism. Mutations or modifications in some of these genes contribute to antibiotic resistance (e.g., *rpoB*, *fusA*, *rpsL*). The top 13 genes in *S. aureus* MRSA encode essential components for vital cellular processes, making them critical for bacterial survival and potential targets for antimicrobial strategies these genes depicted in supplementary file S2. Hamzah et al. (2024) identified key antibiotic resistance genes and chromosomal point mutations, including fusA (fusidic acid), dfrB (trimethoprim), gyrA/grlA (fluoroquinolones) and rpoB (rifampin), as well as genes encoding multidrug efflux pumps such as sepA, lmrS, norA, sdrM, norC, qacA, and mepA. These findings highlight the diverse mechanisms contributing to antibiotic resistance in *S. aureus*.

**Figure 11.**
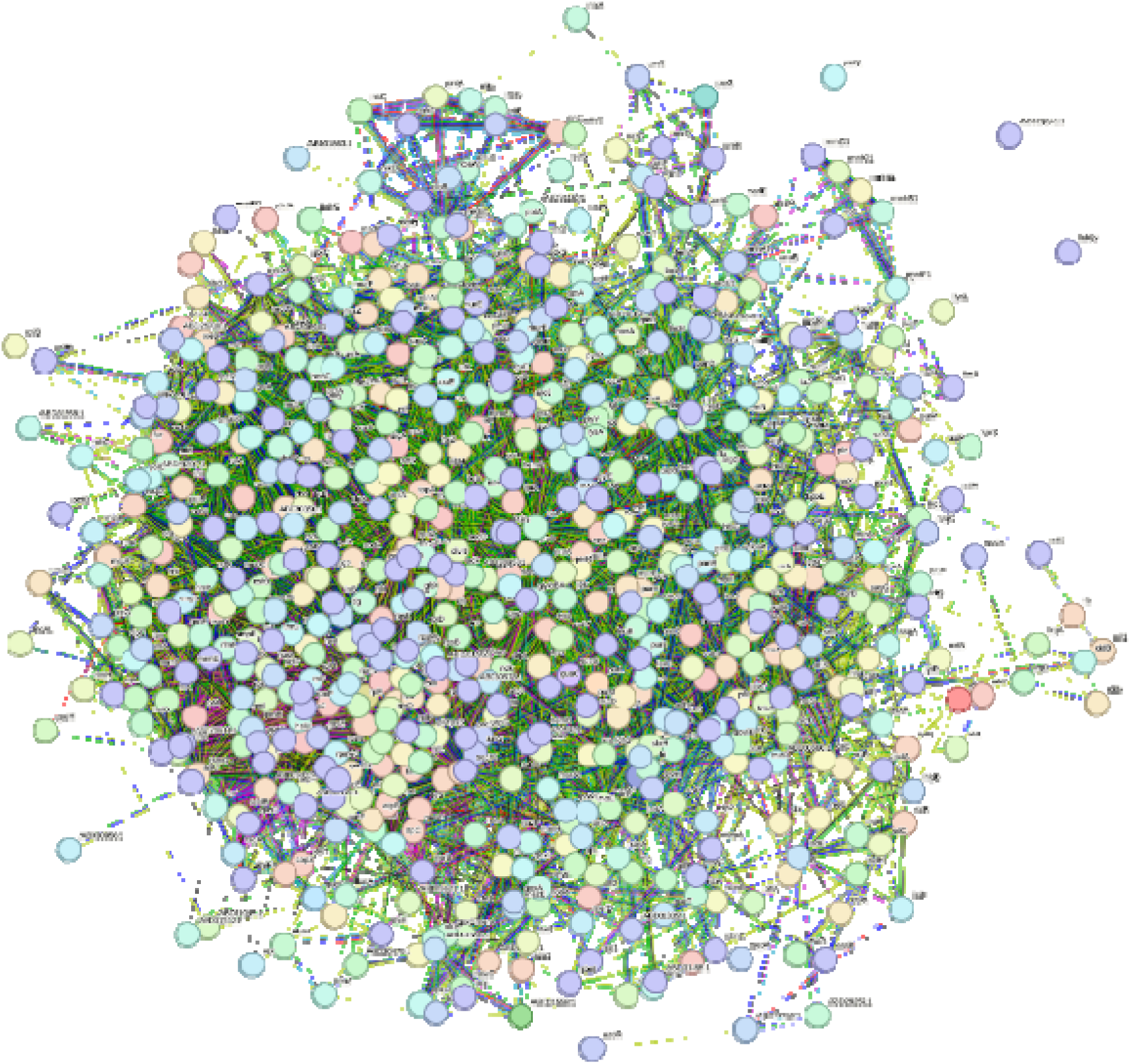
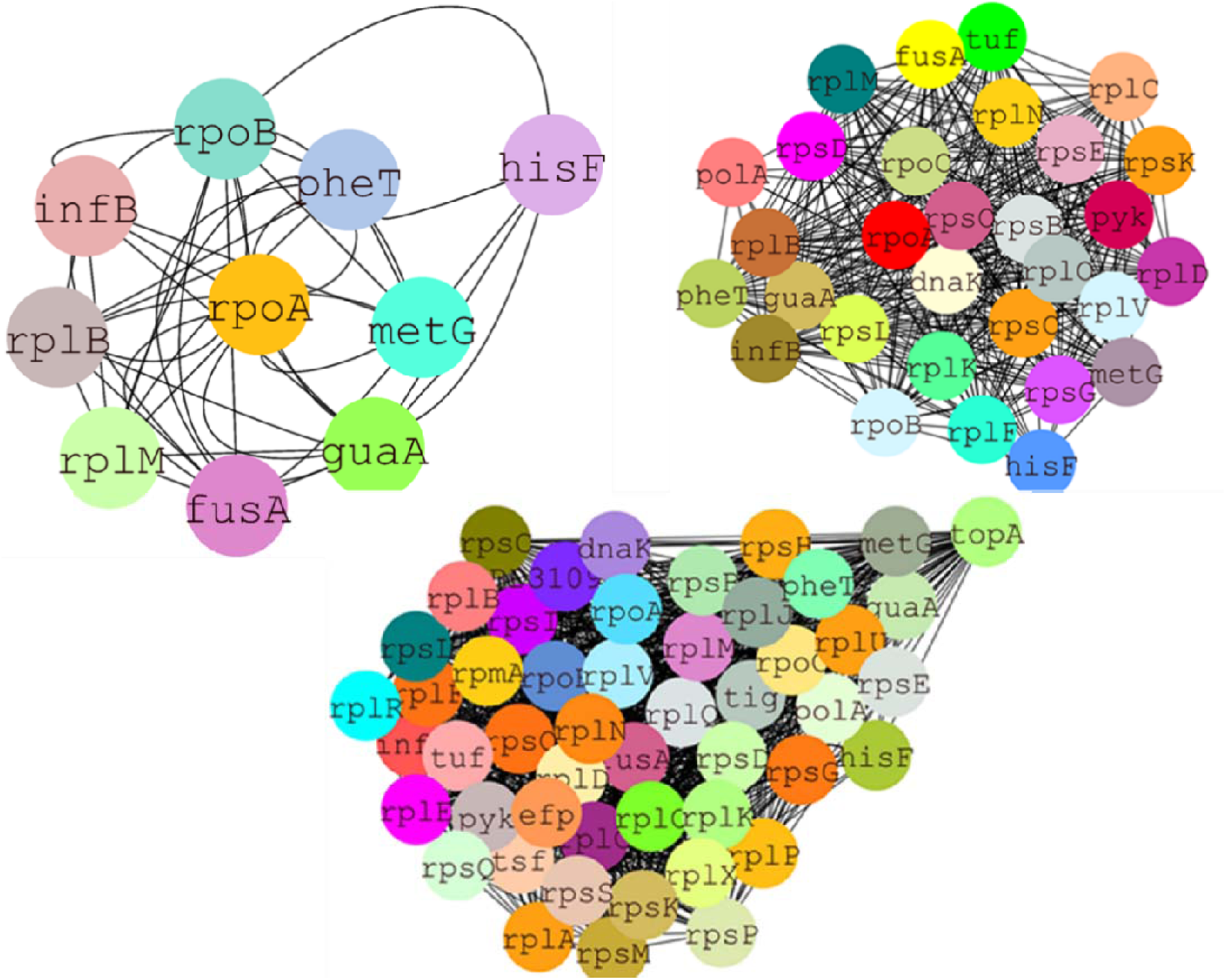
**a.** The STRING network of MRSA strain d1418m22 illustrates the interactions between resistance and biofilm-related genes, with node size corresponding to the number of connections and edge thickness indicating the strength of the correlations. To further explore the relationships, interaction networks were employed to visualize the connections between gene products and highlight those with greater connectivity compared to others. **b, c & d.** Network analysis of the MRSA strain d1418m22 genome, performed using Cytoscape, shows interactions among the top ten, thirty, and fifty resistance genes. Node size reflects the number of connections, while edge thickness indicates correlation strength.

Additionally, several genes in the *MRSA strain* d1418m22 strain were associated with virulence, resistance, and MGEs. Figure 12a illustrates the VFs and their corresponding protein functions, which contribute to the bacterial strain’s virulence mechanisms. The listed genes represent various VFs in MRSA. These genes encode proteins that allow MRSA to evade the human immunity system, establish infections, and impairment host tissues. The *sak* (Staphylokinase) gene activates plasminogen to plasmin, which degrades fibrin clots, allowing MRSA to escape containment and promote bacterial spread within host tissues. Additionally, the hemolysin gamma components *hlgA*, *hlgB*, and *hlgC* play a key role in forming pores that lyse red and white blood cells, contributing to tissue damage and immune evasion. MRSA employs a variety of VFs that enhance its ability to evade the immune system, sustain infections, and damage host tissues. By lysing red blood cells, MRSA obtains nutrients, while leukocyte destruction facilitates immune evasion. The *lukF-PV* and *lukS-PV* genes, encoding PVL components, produce a pore-forming toxin targeting neutrophils. This results in immune cell lysis, a heightened inflammatory response, and severe skin and soft tissue infections. The enterotoxin genes (seo, sei, sel, seg, sem, sen, sec and seu) function as superantigens, inducing T-cell activation and cytokine storms. These genes are implicated in the pathogenesis of staphylococcal food poisoning and toxic shock syndrome (TSS). Collectively, these VFs enable MRSA to effectively infect, persist within the host, and cause significant tissue damage, posing a serious public health challenge. A study by Hamzah et al. (2024) identified 26 virulence genes in 88 MRSA isolates using VirulenceFinder, including host immunity evasion genes (sak, scn), exoenzyme genes (aur, splABE), and toxin genes (edinA, hlgABC, lukED, lukFS, sek–seq, sea–sec, seg–sei, and seu). Most isolates (96.6%, 85/88) carried more than 10 virulence genes. Prevalent genes such as aur, sak, scn, hlgABC, and enterotoxins (egc cluster) were common among MRSA strains from HSNZ and other Malaysian isolates, highlighting their virulence potential. Using the abritAMR tool, 32 virulence determinants were identified, with six consistently found across all assemblies. These include lmrS (multidrug efflux pump) (Floyd et al., 2010), aur (aureolysin, a protease) (Pietrocola et al., 2017), δ- and γ-hemolysins (immune cell lysis) (Verdon et al., 2009), and icaC (cellular adhesion and biofilm formation) (Cramton et al., 1999). Over half of the isolates also carried can (collagen-binding adhesin), promoting bacterial adherence to collagen-rich tissues (Madani et al., 2017). These findings highlight the extensive repertoire of VFs in the studied isolates. In figure 12 b & c network illustrated resistant genes and their mechanism and also mobile genetics elements and their major MobileOG Category showed in network which play significant role in *S. aureus* strain to resistance. The genes listed contribute to antibiotic resistance in *Staphylococcus aureus*, particularly in MRSA. These genes encode proteins that enable the bacterium to resist the effects of various antibiotics, compromising treatment efficacy. Here’s how each gene contributes to resistance and impacts the human host. The mgrA (Multiple Gene Regulator A), A global regulator controlling the expression of efflux pumps and other resistance mechanisms. Enhances MRSA’s ability to expel antibiotics like fluoroquinolones and resist oxidative stress, leading to persistent infections. The genes norA, sdrM, sepA, norC, and LmrS encode efflux pumps that facilitate the expulsion of antibiotics from bacterial cells, leading to decreased intracellular drug levels and promoting antibiotic resistance. kdpD gene encodes a sensor kinase involved in osmoregulation and increases bacterial survival in stressful environments. PC1 Beta-lactamase (blaZ), encodes a β-lactamase enzyme that abolishes β-lactam-antibiotics like penicillin. Gene mecR1 and mecA, mecR1: a sensor and regulator of mecA expression. Confers methicillin resistance, a hallmark of MRSA, making β-lactam antibiotics ineffective. Similarly, in MRSA bacteria, chromosomal point mutations were identified in genes such as *gyr-A*/*grl-A* (fluoroquinolone resistance), *fus-A* (fusidic acid resistance), *dfrB* (trimethoprim resistance), and *rpo-B* (rifampin resistance). Additionally, genes encoding multidrug-efflux-pumps, including, *sepA, mepA*, *norC*, *qacA*, *norA, sdrM*, and *lmrS*, were also detected, highlighting mechanisms contributing to antimicrobial resistance (Hamzah et al., 2024). Multidrug resistance (MDR) efflux pump genes, including *norA*, *LmrS*, *mepA*, and *mepR*, were detected in all *S. aureus* isolates in this research. However, not all of these genes were linked to express phenotypic drug resistance. Notably, other research has similarly stated a high occurrence of these elements in *Staphylococcus* species (Antiabong et al., 2017). Our study highlights the pivotal role of MGEs—including plasmids, transposons, and integrons—in the acquisition, transfer, and regulation of antibiotic resistance genes in MRSA. These elements facilitate the persistence and evolution of resistance traits by contributing to DNA repair, replication, recombination, transcriptional regulation, and stress responses. As depicted in Figure 12c, 80 MGEs were associated with replication, recombination, and repair within the mobileOG category. Additionally, 64 were linked to integration/excision, 48 to phage activity, 45 to gene transfer, and 18 to functions encompassing stability, transfer, and defense mechanisms.

**Figure 12.**
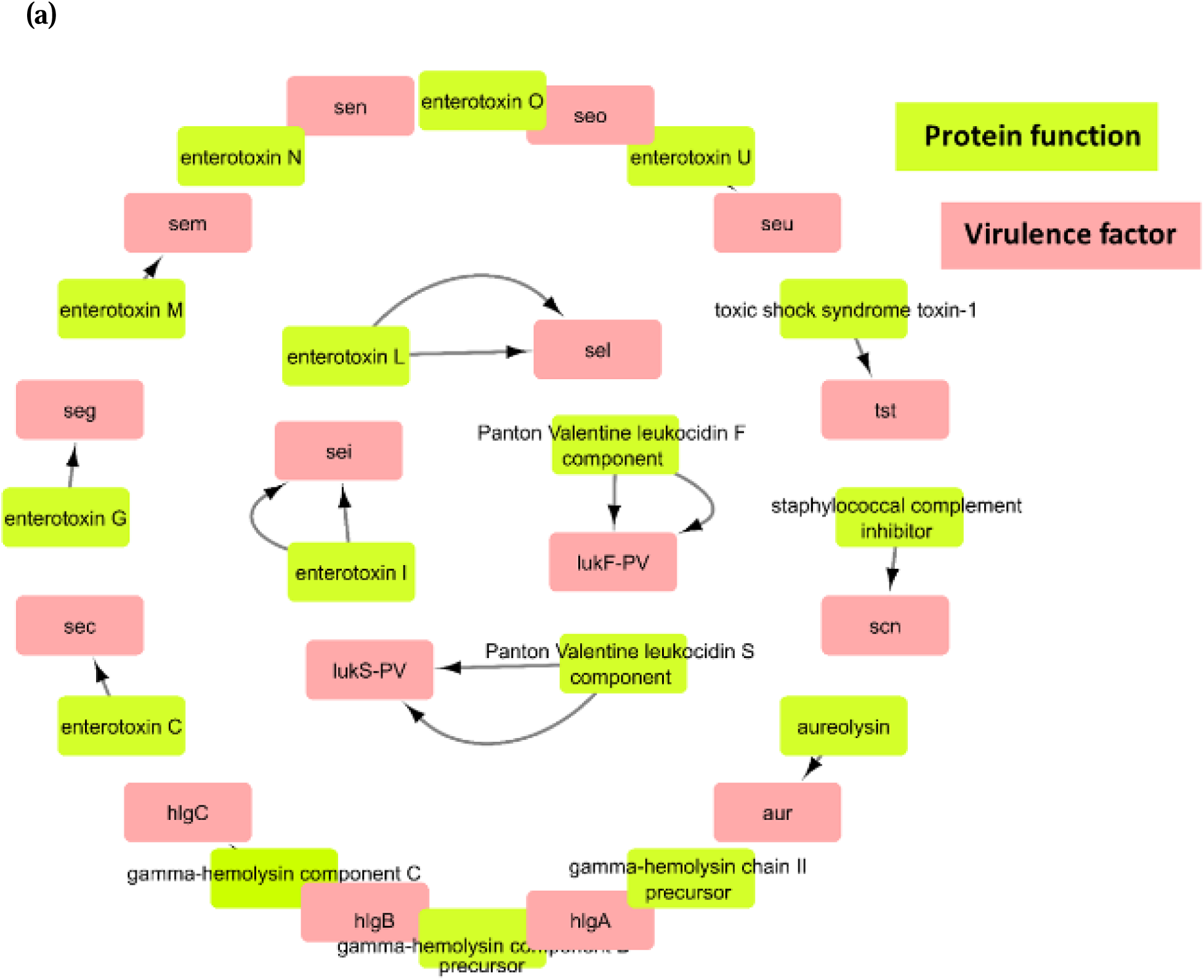
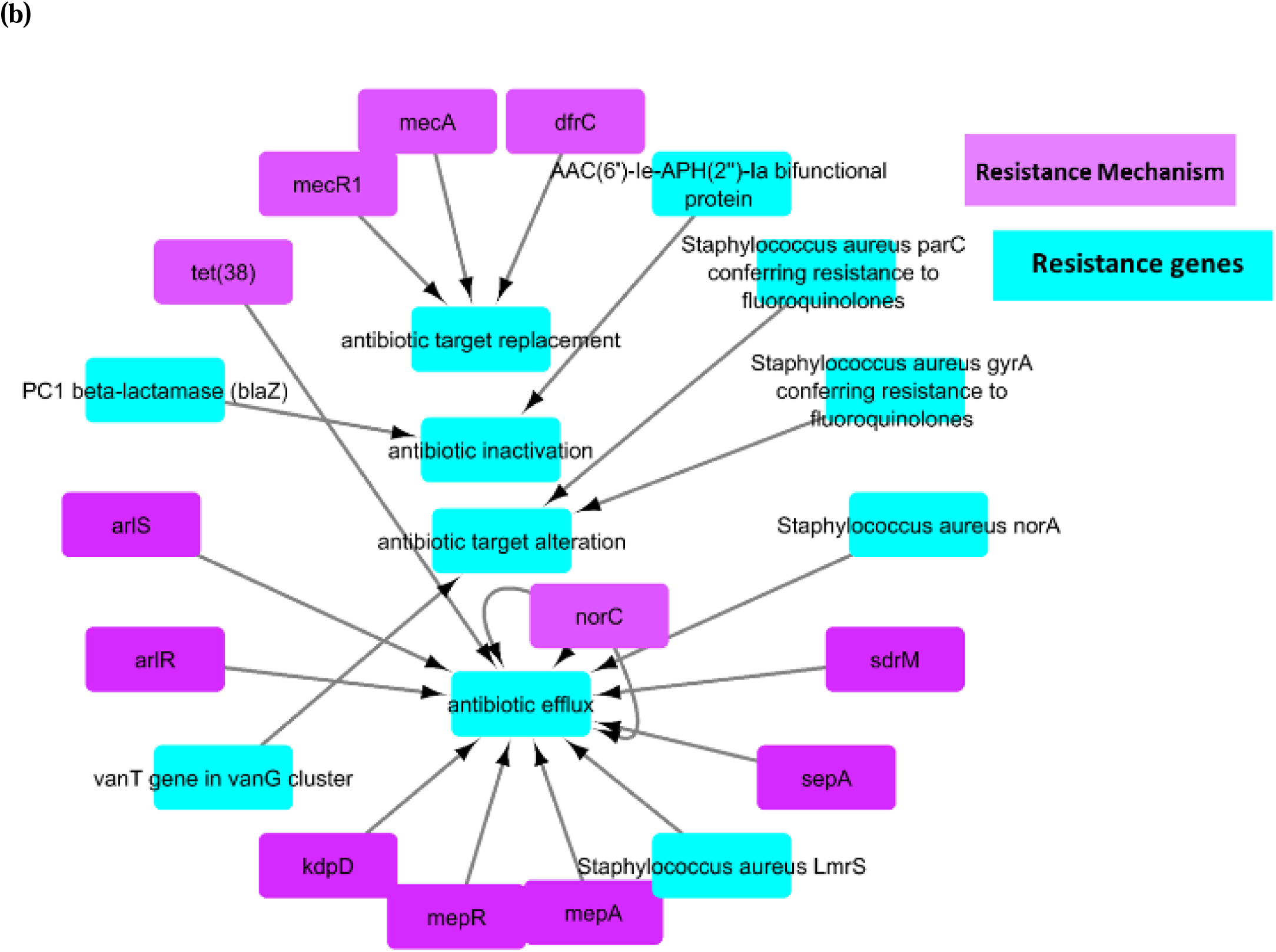
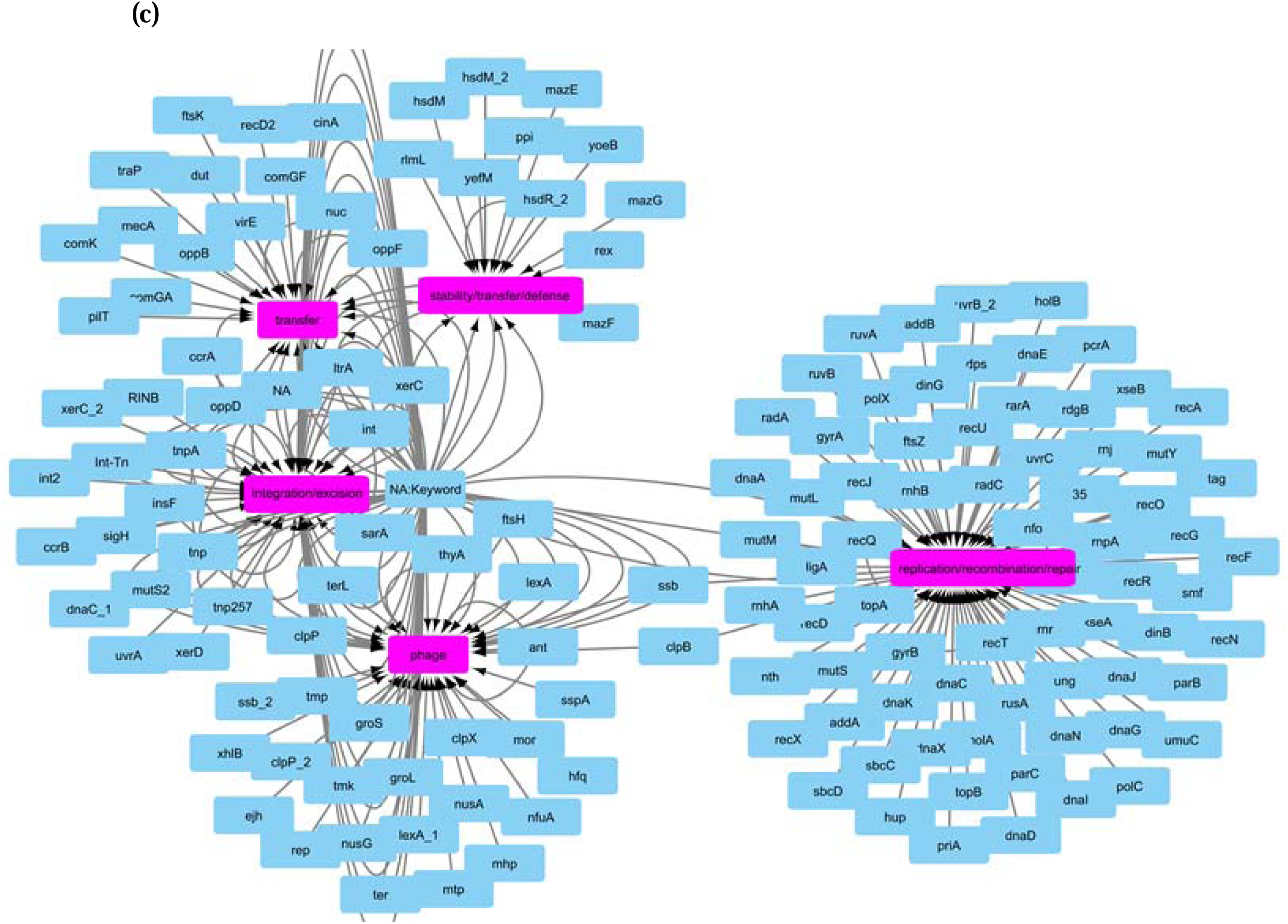
a, b &. **c.** (a) Network analysis of virulence genes and their interactions with protein mechanisms. (b) Resistance gene networks and the functional mechanisms of these genes within the bacterial strain. (c) MGEs categorized by their primary mobileOG classifications.

## Conclusion

This study provides a comprehensive genomic characterization of MRSA strain d1418m22. WGS analysis revealed a genome size of 2.78 Mb, harboring 2,625 predicted genes. Functional annotation identified genes involved in various cellular processes, including metabolism, virulence, and antibiotic resistance. The occurrence of SCCmec-type-IVa, MGEs, and VFs, such as PVL and enterotoxins, highlights the strain’s potential for pathogenicity and its contribution to the spread of antibiotic resistance. Interaction network analysis identified key hub genes involved in essential cellular functions. These findings deliver valuable understandings into the genetic makeup of MRSA strain d1418m22, aiding in the progress of effective strategies for prevention and control of MRSA infections.

## AUTHOR CONTRIBUTIONS

Daraksha Iram: Conceptualization, Visualization, Writing – original draft, Writing – review | Manish Singh Sansi: Conceptualization, Visualization, Writing – original draft, Writing – review | Satpal Dixit: Writing – original draft, Writing – review and editing.

## DATA AVAILABILITY

The research described in this article did not involve the use of any original data.

## References

1. Abdulgader, S.M.; Shittu, A.O.; Nicol, M.P.; Kaba, M. Molecular Epidemiology of Methicillin-Resistant *Staphylococcus aureus* in Africa: A Systematic Review. Front. Microbiol. 2015, 6, 348.

2. Abdulgader, S.M.; Shittu, A.O.; Nicol, M.P.; Kaba, M. Molecular Epidemiology of Methicillin-Resistant *Staphylococcus aureus* in Africa: A Systematic Review. Front. Microbiol. 2015, 6, 348.

3. Alcock, et al. 2023. CARD 2023: expanded curation, support for machine learning, and resistome prediction at the Comprehensive Antibiotic Resistance Database. Nucleic Acids Research, 51(D1): D690–D699

4. Ali, M. S., Isa, N. M., Abedelrhman, F. M., Alyas, T. B., Mohammed, S. E., Ahmed, A. E., & Mohamed, S. B. (2019). Genomic analysis of *methicillin-resistant Staphylococcus aureus* strain SO-1977 from Sudan. BMC microbiology, 19, 1–9.

5. Arndt, D.; Grant, J.R.; Marcu, A.; Sajed, T.; Pon, A.; Liang, Y.; Wishart, D.S. PHASTER: A Better, Faster Version of the PHAST Phage Search Tool. Nucleic Acids Res. 2016, 44, W16–W21.

6. Asadollahi P, Farahani NN, Mirzaii M, Khoramrooz SS, van Belkum A, Asadollahi K, Dadashi M, Darban-Sarokhalil D. 2018. Distribution of the most prevalent spa types among clinical isolates of methicillin-resistant and -susceptible Staphylococcus aureus around the world: a review. Front Microbiol 9:163.

7. Atshan S.S., Shamsudin M.N., Karunanidhi A., van Belkum A., Lung L.T.T., Sekawi Z., Nathan J.J., Ling K.H., Seng J.S.C., Ali A.M., et al. Quantitative PCR analysis of genes expressed during biofilm development of methicillin resistant Staphylococcus aureus (MRSA) Infect. Genet. Evol. 2013;18: 106–112.

8. Baig, S.; Johannesen, T.B.; Overballe-Petersen, S.; Larsen, J.; Larsen, A.R.; Stegger, M. Novel SCC*mec* Type XIII (9A) identified in an ST152 Methicillin-Resistant *Staphylococcus aureus*. Infect. Genet. Evol. 2018, 61, 74–76.

9. Becker K, Heilmann C, Peters G. 2014. Coagulase-negative staphylococci. Clin Microbiol Rev 27:870–926.

10. Ben Nejma, M.; Mastouri, M.; Bel Hadj Jrad, B.; Nour, M. Characterization of ST80 Panton-Valentine Leukocidin-positive community-acquired methicillin resistant *Staphylococcus aureus* clone in Tunisia. Diagn. Microbiol. Infect. Dis. 2013, 77, 20–24.

11. Besser JM, Carleton HA, Trees E, Stroika SG, Hise K, Wise M, Gerner-Smidt P. 2019. Interpretation of whole-genome sequencing for enteric disease surveillance and outbreak investigation. Foodborne Pathog Dis 16: 504– 512.

12. Bortolaia, V.; Kaas, R.F.; Ruppe, E.; Roberts, M.C.; Schwarz, S.; Cattoir, V.; Philippon, A.; Allesoe, R.L.; Rebelo, A.R.; Florensa, A.F.;, et al. ResFinder 4.0 for predictions of phenotypes from genotypes. J. Antimicrob. Chemother. 2020, 75, 3491–3500.

13. Bouchami, O.; Achour, A.; Ben Hassan, A. Typing of staphylococcal cassette chromosome *mec* encoding methicillin resistance in *Staphylococcus aureus* strains isolated at the bone marrow transplant centre of Tunisia. Curr. Microbiol. 2009, 59, 380–385.

14. Brown CL, Mullet J, Hindi F, Stoll JE, et al. mobileOG-db: a Manually Curated Database of Protein Families Mediating the Life Cycle of Bacterial Mobile Genetic Elements. Appl Environ Microbial. 2022 Sep 22; 88(18).

15. Carattoli, A.; Hasman, H. PlasmidFinder and In Silico PMLST: Identification and Typing of Plasmid Replicons in Whole-Genome Sequencing (WGS). Methods Mol. Biol. 2020, 2075, 285–294.

16. Carattoli, A.; Zankari, E.; Garciá-Fernández, A.; Larsen, M.V.; Lund, O.; Villa, L.; Aarestrup, F.M.; Hasman, H. In Silico Detection and Typing of Plasmids Using Plasmidfinder and Plasmid Multilocus Sequence Typing. Antimicrob. Agents Chemother. 2014, 58, 3895–3903.

17. Carattoli, A.; Zankari, E.; Garcia-Fernandez, A.; Voldby Larsen, M.; Lund, O.; Villa, L.; Møller Aarestrup, F.; Hasman, H. PlasmidFinder and pMLST: In Silico detection and typing of plasmids. Antimicrob. Agents Chemother. 2014, 58, 3895–3903.

18. Cerca, N.; Brooks, J. L.; Jefferson, K. K. Regulation of the intercellular adhesin locus regulator (icaR) by SarA, sigmaB, and IcaR in Staphylococcus aureus. J. Bacteriol. 2008, 190 (19), 6530−3

19. Chambers HF, DeLeo FR. 2009. Waves of resistance: staphylococcus aureus in the antibiotic era. Nat Rev Microbiol 7:629–641.

20. Chen, L.; Xiong, Z.; Sun, L.; Yang, J.; Jin, Q. VFDB 2012 Update: Toward the Genetic Diversity and Molecular Evolution of Bacterial Virulence Factors. Nucleic Acids Res. 2012, *40*, 641–645.

21. Chin, C. H.; Chen, S. H.; Wu, H. H.; Ho, C. W.; Ko, M. T.; Lin, C. Y. cytoHubba: identifying hub objects and sub-networks from complex interactome. BMC Syst. Biol. 2014, 8 (Suppl 4), S11.

22. 22. Clausen, P.T.L.C.; Aarestrup, F.M.; Lund, O. Rapid and precise alignment of raw reads against redundant databases with KMA. BMC Bioinform. 2018, 19, 307.

23. CLSI (2023). “Performance standards for antimicrobial disk and dilution susceptibility tests for Bacteria isolated from animals” in CLSI Document VET01-S. 6th ed. Eds. C. R. Burbick (Pittsburgh, PA, USA: CLSI)

24. Couvin D, Bernheim A, Toffano-Nioche C, Touchon M. et al. CRISPRCasFinder, an update of CRISRFinder, includes a portable version, enhanced performance and integrates search for Cas proteins. Nucleic Acids Res. 2018 Jul 2; 46(W1): W246–W251.

25. Cramton, S.E., Gerke, C., Schnell, N.F., Nichols, W.W., Gotz, F., 1999. The intercellular adhesion (ica) locus is present in Staphylococcus aureus and is required for biofilm formation. Infect. Immun. 67 (10), 5427–5433.

26. Cue, D., Lei, M. G., and Lee, C. Y. (2012). Genetic regulation of the intercellular adhesion locus in staphylococci. Front. Cell. Infect. Microbiol. 2, 38.

27. Dhar, H.; Al-Busaidi, I.; Rathi, B.; Nimre, E.A.; Sachdeva, V.; Hamdi, I. A study of post-caesarean section wound infections in a regional referral hospital, Oman. Sultan Qaboos Univ. Med. J. 2014, 14, 211.

28. Elhawy, M. I., Huc-Brandt, S., Pätzold, L., Gannoun-Zaki, L., Abdrabou, A. M. M., Bischoff M., et al. (2021). Th Phosphoarginine phosphatase Ptp B from *Staphylococcus aureus* is involved in bacterial stress adaptation during infection. Cells 10:645.

29. Eyre DW, Golubchik T, Gordon NC, Bowden R, Piazza P, Batty EM, Ip CLC, Wilson DJ, Didelot X, O’Connor L, Lay R, Buck D, Kearns AM, Shaw A, Paul J, Wilcox MH, Donnelly PJ, Peto TEA, Walker AS, Crook DW. 2012. A pilot study of rapid benchtop sequencing of Staphylococcus aureus and Clostridium difficile for outbreak detection and surveillance. BMJ Open 2: e001124.

30. Floyd, J.L., Smith, K.P., Kumar, S.H., Floyd, J.T., Varela, M.F., 2010. LmrS is a multidrug efflux pump of the major facilitator superfamily from Staphylococcus aureus. Antimicrob. Agents Chemother. 54 (12), 5406–5412.

31. Froggatt JW, Johnston JL, Galetto DW, Archer GL. 1989. Antimicrobial resistance in nosocomial isolates of Staphylococcus haemolyticus. Antimicrob Agents Chemother 33:460–466.

32. Funaki T, Yasuhara T, Kugawa S, Yamazaki Y, Sugano E, Nagakura Y, Yoshida K, Fukuchi K. 2019. SCCmec typing of PVL-positive community acquired Staphylococcus aureus (CA-MRSA) at a Japanese hospital. Heliyon 5: e01415.

33. Gelalcha, B. D., Agga, G. E., and Dego, O. K. (2022). “Antimicrobial usage for the management of mastitis in the USA: impacts on antimicrobial resistance and potential alternative approaches” in Mastitis in dairy cattle, sheep and goats, Eds. O. K. Dego (Intech Open) 1–21.

34. Gillet, Y.; Issartel, B.; Vanhems, P.; Fournet, J.-C.; Lina, G.; Bes, M.; Vandenesch, F.; Piémont, Y.; Brousse, N.; Floret, D.;, et al. Association between *Staphylococcus aureus* strains carrying gene for Panton-Valentine Leukocidin and highly lethal necrotizing pneumonia in young immunocompetent patients. Lancet 2002, 359, 753–759.

35. Grant, J.R.; Stothard, P. The CGView server: A comparative genomics tool for circular genomes. Nucleic Acids Res. 2008, 36, W181–W184.

36. Grant, J.R.; Stothard, P. The CGView server: A comparative genomics tool for circular genomes. Nucleic Acids Res. 2008, 36, W181–W184.

37. Guo, Y., Song, G., Sun, M., Wang, J., and Wang, Y. (2020). Prevalence and therapies of antibiotic-resistance in *Staphylococcus aureus*. Front. Cell. Infect. Microbiol. 10, 107.

38. Harris SR, Cartwright EJP, Török ME, Holden MTG, Brown NM, OgilvyStuart AL, Ellington MJ, Quail MA, Bentley SD, Parkhill J, Peacock SJ. 2013. Whole-genome sequencing for analysis of an outbreak of meticillin-resistant Staphylococcus aureus: a descriptive study. Lancet Infect Dis 13: 130–136.

39. Holtfreter, S.; Grumann, D.; Schmudde, M.; Nguyen, H.T.T.; Eichler, P.; Strommenger, B.; Kopron, K.; Kolata, J.; Giedrys-Kalemba, S.; Steinmetz, I.;, et al. Clonal distribution of superantigen genes in clinical *Staphylococcus aureus* isolates. J. Clin. Microbiol. 2007,45, 2669–2680.

40. Ikonomidis, A.; Vasdeki, A.; Kristo, I.; Maniatis, A.N.; Tsakris, A.; Malizos, K.N.; Pournaras, S. Association of Biofilm Formation and Methicillin-Resistance with Accessory Gene Regulator (*Agr*) Loci in Greek *Staphylococcus aureus* Clones. Microb. Pathog. 2009, 47, 341–344.

41. Iram, D., Sansi, M. S., Puniya, A. K., Gandhi, K., Meena, S., & Vij, S. (2024). Phenotypic and molecular characterization of clinically isolated antibiotics-resistant *S. aureus* (MRSA), *E. coli* (ESBL) and *Acinetobacter* 1379 bacterial strains. Brazilian Journal of Microbiology, 1–20.

42. Jamrozy, D., Coll, F., Mather, A. E., Harris, S. R., Harrison, E. M., MacGowan, A., & Peacock, S. J. (2017). Evolution of mobile genetic element composition in an epidemic methicillin-resistant Staphylococcus aureus: temporal changes correlated with frequent loss and gain events. BMC genomics, 18, 1–12.

43. Jemili-Ben Jomaa, M.; Boutiba-Ben Boubaker, I.; Ben Redjeb, S. Identification of staphylococcal cassette chromosome *mec* encoding methicillin resistance in *Staphylococcus aureus* isolates at Charles Nicolle Hospital of Tunis. Pathol. Biol. 2006, 54, 453–455.

44. Jia, B.; Raphenya, A.R.; Alcock, B.; Waglechner, N.; Guo, P.; Tsang, K.K.; Lago, B.A.; Dave, B.M.; Pereira, S.; Sharma, A.N.;, et al. CARD 2017: Expansion and Model-Centric Curation of the Comprehensive Antibiotic Resistance Database. Nucleic Acids Res. 2017, 45, D566–D573.

45. Johansson, M.H.K.; Bortolaia, V.; Tansirichaiya, S.; Aarestrup, F.M.; Roberts, A.P.; Petersen, T.N. Detection of mobile genetic elements associated with antibiotic resistance in *Salmonella enterica* using a newly developed web tool: Mobile Element Finder. J. Antimicrob. Chemother. 2020, 76, 101–109.

46. Kechrid, A.; P érez-V ázquez, M.; Smaoui, H.; Hariga, D.; Rodr íguez-Baños, M.; Vindel, A.; Baquero, F.; Cantón, R.; del Campo, R. Molecular Analysis of Community-Acquired Methicillin-Susceptible and Resistant *Staphylococcus aureus* Isolates Recovered from Bacteraemic and Osteomyelitis Infections in Children from Tunisia. Clin. Microbiol. Infect. 2011, 17, 1020–1026.

47. Khasapane, N. G., Koos, M., Nkhebenyane, S. J., Khumalo, Z. T., Ramatla, T., and Thkisoe, O. (2024). Detection of Staphylococcus isolates and their antimicrobial resistance profies and virulence genes from subclinical mastitis cattle Milk using MALDI-TOF MS, PCR and sequencing in Free State Province, South Africa. Animals 14:154.

48. Khosravi AD, Jenabi A. Montazeri EA. Distribution of genes encoding resistance to aminoglycoside modifying enzymes in methicillinresistant *Staphylococcus aureus* (MRSA) strains. Kaoh Jour Med Scie. 2017;33(12):587–93

49. Klibi, A., Jouini, A., Gómez, P., Slimene, K., Ceballos, S., Torres, C., et al. (2018). Molecular characterization and clonal diversity of methicillin-resistant and-susceptible *Staphylococcus aureus isolates* of milk of cows with clinical mastitis in Tunisia. Microb. Drug Resist. 24, 1210–1216.

50. Kmiha, S., Jouini, A., Zerriaa, N., Hamrouni, S., Thabet, L., & Maaroufi, A. (2023). Methicillin-Resistant Staphylococcus aureus Strains Isolated from Burned Patients in a Tunisian Hospital: Molecular Typing, Virulence Genes, and Antimicrobial Resistance. Antibiotics, 12(6), 1030.

51. Lakhundi, S.; Zhang, K. Methicillin-Resistant *Staphylococcus aureus*: Molecular Characterization, Evolution, and Epidemiology. Clin. Microbiol. Rev. 2018, 31, e00020–18.

52. Li, L. (2008). Complete genome sequence and strain diffrentiation of Mycobacterium avium subspecies paratuberculosis. Minneapolis, United States: University of Minnesota.

53. Liao, F., Mo, Z., Gu, W., Xu, W., Fu, X., & Zhang, Y. (2020). A comparative genomic analysis between *methicillin-resistant Staphylococcus aureus* strains of hospital acquired and community infections in Yunnan province of China. BMC infectious diseases, 20, 1–12.

54. Lina, G.; Piemont, Y.; Godail-Gamot, F.; Bes, M.; Peter, M.-O.; Gauduchon, V.; Vandenesch, F.; Etienne, J. Involvement of Panton-Valentine Leukocidin producing *Staphylococcus aureus* in primary skin infections and pneumonia. Clin. Infect. Dis. 1999, 29, 1128–1132

55. Lindsay JA. 2014. Evolution of Staphylococcus aureus and MRSA during outbreaks. Infect Genet Evol 21:548–553.

56. Lindsay, J. A., Ruzin, A., Ross, H. F., Kurepina, N., & Novick, R. P. (1998). The gene for toxic shock toxin is carried by a family of mobile pathogenicity islands in Staphylococcus aureus. Molecular microbiology, 29(2), 527–543.

57. Liu, B.; Zheng, D.; Jin, Q.; Chen, L.; Yang, J. VFDB 2019: A Comparative Pathogenomic Platform with an Interactive Web Interface. Nucleic Acids Res. 2019, *47*, D687–D692.

58. Liu, K., Tao, L., Li, J., Fang, L., Cui, L., Li, J., et al. (2020). Characterization of *Staphylococcus aureus* isolates from cases of clinical bovine mastitis on large-scale Chinese dairy farms. Front Vet Sci 7:580129.

59. Livermore, D. M. (2000). Antibiotic resistance in staphylococci. Int. J. Antimicrob. Agents 16, 3–10.

60. Maarouf L, Omar H, El-Nakeeb M, Abouelfetouh A. 2020. Prevalence and mechanisms of linezolid resistance among staphylococcal clinical isolates from Egypt. Eur J Clin Microbiol Infect Dis 40:815–823

61. Madani, A., Garakani, K., Mofrad, M.R.K., 2017. Molecular mechanics of *Staphylococcus aureus* adhesin, CNA, and the inhibition of bacterial adhesion by stretching collagen. PLoS One 12 (6), e0179601.

62. Magill, S.S.; Hellinger, W.; Cohen, J.; Kay, R.; Bailey, C.; Boland, B.; Carey, D.; de Guzman, J.; Dominguez, K.; Edwards, J.;, et al. Prevalence of healthcare-associated infections in acute care hospitals in Jacksonville, Florida. Infect. Control Hosp. Epidemiol. 2012, 33, 283–291.

63. Maity, S., Das, D., and Ambatipudi, K. (2020). Quantitative alterations in bovine milk proteome from healthy, subclinical and clinical mastitis during S. aureus infection. J. Proteom. 223, p.103815.

64. Maity, S., Das, D., and Ambatipudi, K. (2020). Quantitative alterations in bovine milk proteome from healthy, subclinical and clinical mastitis during S. aureus infection. J. Proteom. 223, p.103815.

65. Mottola, C.; Semedo-Lemsaddek, T.; Mendes, J.J.; Melo-Cristino, J.; Tavares, L.; Cavaco-Silva, P.; Oliveira, M. Molecular typing, virulence traits and antimicrobial resistance of diabetic foot staphylococci. J. Biomed. Sci. 2016, 23, 33.

66. Mpogoro, F.J.; Mshana, S.E.; Mirambo, M.M.; Kidenya, B.R.; Gumodoka, B.; Imirzalioglu, C. Incidence and predictors of surgical site infections following caesarean sections at Bugando Medical Centre, Mwanza, Tanzania. Antimicrob. Resist. Infect. Control 2014, 3, 25.

67. Naorem, R. S., Blom, J., & Fekete, C. (2021). Genome-wide comparison of four MRSA clinical isolates from Germany and Hungary. PeerJ, 9, e10185.

68. Naushad, S., Naqvi, S. A., Nobrega, D., Luby, C., Kastelic, J. P., Barkema, H. W., et al. (2019). Comprehensive virulence gene profiing of bovine non-aureus staphylococci based on whole-genome sequencing data. Msystems 4, 10–1128.

69. Nourbakhsh F., Namvar A.E. Detection of genes involved in biofilm formation in Staphylococcus aureus isolates. GMS Hyg. Infect. Control. 2016;11: Doc07.

70. Oliveira DC, de Lencastre H. Methicillin-resistance in Staphylococcus aureus is not affected by the overexpression in trans of the mecA gene repressor: a surprising observation. PLoS One. 2011;6(8): e23287.

71. Omwenga, I., Aboge, G. O., Mitema, E. S., Obiero, G., Ngaywa, C., Ngwili, N., et al. (2021). Antimicrobial usage and detection of multidrug-resistant *Staphylococcus aureus*, including methicillin-resistant strains in raw milk of livestock from northern Kenya. Microb. Drug Resist. 27, 843–854.

72. Pattabhiramaiah, M., and Mallikarjunaiah, S. (2023). “Foodborne Pathogens and food safety regulations” in Food microbial and molecular biology (Florida, USA: Apple Academic Press), 179–212.

73. Pietrocola, G., Nobile, G., Rindi, S., Speziale, P., 2017. Staphylococcus aureus manipulates innate immunity through own and host-expressed proteases. Front. Cell. Infect. Microbiol. 7, 166.

74. Pillai, S.K., Sakoulas, G., Wennersten, C., Eliopoulos, G.M., Moellering Jr., R.C., Ferraro, M.J., Gold, H.S., 2002. Linezolid resistance in Staphylococcus aureus: characterization and stability of resistant phenotype. J. Infect. Dis. 186 (11), 1603–1607.

75. Pobiega M, Wójkowska-Mach J, Heczko PB. 2013. Typing of Staphylococcus aureus in order to determine the spread of drug-resistant strains inside and outside hospital environment. Przegl Epidemiol 67:435–438

76. Sabri I, Adwan K, Essawi TA, Farraj MA. 2013. Molecular characterization of methicillin-resistant Staphylococcus aureus isolates in three different Arab world countries. Eur J Microbiol Immunol (Bp) 3:183–187.

77. Schmitz, F.-J., Jones Mark, E., Hofmann, B., Hansen, B., Scheuring, S., Lückefahr, M., Kohrer, K., 1998. Characterization of grlA, grlB, gyrA, and gyrB mutations in 116 unrelated isolates of Staphylococcus aureus and effects of mutations on ciprofloxacin MIC. Antimicrob. Agents Chemother. 42 (5), 1249–1252.

78. Seemann T, “Prokka: Rapid Prokaryotic Genome Annotation”, Bioinformatics, 2014 Jul 15;30(14):2068–9

79. Shannon, P.; Markiel, A.; Ozier, O.; Baliga, N. S.; Wang, J. T.; Ramage, D.; Amin, N.; Schwikowski, B.; Ideker, T. Cytoscape: a software environment for integrated models of biomolecular interaction networks. Genome Res. 2003, 13 (11), 2498−504.

80. 80. Sivakumar, R., Pranav, P. S., Annamanedi, M., Chandrapriya, S., Isloor, S., Rajendhran, J., et al. (2023). Genome sequencing and comparative genomic analysis of bovine mastitis-associated *Staphylococcus aureus* strains from India. BMC Genomics 24:44.

81. Smyth, E.T.; McIlvenny, G.; Enstone, J.E.; Emmerson, A.M.; Humphreys, H.; Fitzpatrick, F.; Davies, E.; Newcombe, R.G.; Spencer, R.C. Hospital Infection Society Prevalence Survey Steering Group. Four country healthcare associated infection prevalence survey 2006: Overview of the results. J. Hosp. Infect. 2008, 69, 230–248.

82. Stefani S, Chung DR, Lindsay JA, Friedrich AW, Kearns AM, Westh H, MacKenzie FM. 2012. Meticillin-resistant Staphylococcus aureus (MRSA): global epidemiology and harmonisation of typing methods. Int J Antimicrob Agents 39:273–282.

83. Steinig EJ, Duchene S, Robinson DA, Monecke S, Yokoyama M, Laabei M, Slickers P, Andersson P, Williamson D, Kearns A, Goering RV, Dickson E, Ehricht R, Ip M, O’sullivan MVN, Coombs GW, Petersen A, Brennan G, Shore AC, Coleman DC, Pantosti A, de Lencastre H, Westh H, Kobayashi N, Heffernan H, Strommenger B, Layer F, Weber S, Aamot HV, Skakni L, Peacock SJ, Sarovich D, Harris S, Parkhill J, Massey RC, Holden MTG, Bentley SD, Tong SYC. 2019. Evolution and global transmission of a multidrug-resistant, community-associated methicillin-resistant staphylococcus aureus lineage from the Indian subcontinent.

84. Struelens MJ. 1996. Consensus guidelines for appropriate use and evaluation of microbial epidemiologic typing systems. Clin Microbiol Infect 2: 2–11.

85. Tokajian S. 2014. New epidemiology of Staphylococcus aureus infections in the Middle East. Clin Microbiol Infect 20:624–628.

86. Tong SYC, Davis JS, Eichenberger E, Holland TL, Fowler VG. 2015. Staphylococcus aureus infections: epidemiology, pathophysiology, clinical manifestations, and management. Clin Microbiol Rev 28:603–661.

87. Ullah, N., Nasir, S., Ishaq, Z., Anwer, F., Raza, T., Rahman, M., & Ali, A. (2022). Comparative Genomic Analysis of a Panton–Valentine Leukocidin-Positive ST22 Community-Acquired Methicillin-Resistant Staphylococcus aureus from Pakistan. Antibiotics, 11(4), 496.

88. 88. Vanderhaeghen, W., Piepers, S., Leroy, F., Van Coillie, E., Haesebrouck, F., and De Vliegher, S. (2015). Identifiation, typing, ecology and epidemiology of coagulase negative staphylococci associated with ruminants. Vet. J. 203, 44–51.

89. Vargová, M., Zigo, F., Výrostková, J., Farkašová, Z., and Rehan, I. F. (2023). Biofimproducing ability of *Staphylococcus aureus* obtained from surfaces and Milk of Mastitic cows. Vet Sci 10:386.

90. Verdon, J., Girardin, N., Lacombe, C., Berjeaud, J.-M., Héchard, Y., 2009. δ-hemolysin, an update on a membrane-interacting peptide. Peptides 30 (4), 817–823.

91. Vernikos GS, Parkhill J. Interpolated variable order motifs for identification of horizontally acquired DNA: revisiting the Salmonella pathogenicity islands. Bioinformatics. 2006 Sep 15; 22(18):2196–203.

92. Viering, B.; Cunningham, T.; King, A.; Blackledge, M. S.; Miller, H. B. Brominated Carbazole with Antibiotic Adjuvant Activity Displays Pleiotropic Effects in MRSA’s Transcriptome. ACS Chem. Biol. 2022, 17, 1239.

93. Víquez-Molina, G.; Aragón-Sánchez, J.; Pérez-Corrales, C.; Murillo-Vargas, C.; López-Valverde, M.E.; Lipsky, B.A. Virulence factor genes in *Staphylococcus aureus* isolated from diabetic foot soft tissue and bone infections. Int. J. Low. Extrem. Wounds 2018,17, 36–41.

94. Xu, Y.; Qian, S.-Y.; Yao, K.-H.; Dong, F.; Song, W.-Q.; Sun, C.; Yang, X.; Zhen, J.-H.; Liu, X.-Q.; Lv, Z.-Y.;, et al. Clinical and molecular characteristics of *Staphylococcus aureus* isolated from chinese children: Association among the *agr* groups and genotypes, Virulence Genes and Disease Types. World J. Pediatr. 2021, 17, 180–188.

95. Yezli S, Shibl AM, Livermore DM, Memish ZA. 2012. Antimicrobial resistance among Gram-positive pathogens in Saudi Arabia. J Chemother 24: 125–136.

96. Zankari, E.; Allesoe, R.; Joensen, K.G.; Cavaco, L.M.; Lund, O.; Aarestrup, F.M. PointFinder: A novel web tool for WGS-based detection of antimicrobial resistance associated with chromosomal point mutations in bacterial pathogens. J. Antimicrob. Chemother. 2017, 72, 2764–2768.

97. Zhou, Y.; Liang, Y.; Lynch, K.H.; Dennis, J.J.; Wishart, D.S. PHAST: A Fast Phage Search Tool. Nucleic Acids Res. 2011, 39, 347–352.

98. Abdulgader, S.M.; Shittu, A.O.; Nicol, M.P.; Kaba, M. Molecular Epidemiology of Methicillin-Resistant Staphylococcus aureus in Africa: A Systematic Review. Front. Microbiol. 2015, 6, 348.

99. O’Hara, F.P.; Suaya, J.A.; Ray, G.T.; Baxter, R.; Brown, M.L.; Mera, R.M.; Close, N.M.; Thomas, E.; Amrine-Madsen, H. Spa typing and multilocus sequence typing show comparable performance in a macroepidemiologic study of *Staphylococcus aureus* in the United States. Microb. Drug Resist. 2016, 22, 88–96.

100. Chikhi R, Medvedev P. 2014. Informed and automated k-mer size selection for genome assembly. Bioinformatics 30:31–37.

101. Zerbino DR, Birney E. 2008. Velvet: algorithms for *de novo* short read assembly using de Bruijn graphs. Genome Res 18:821–829.

102. Bankevich A, Nurk S, Antipov D, Gurevich AA, Dvorkin M, Kulikov AS, Lesin VM, Nikolenko SI, Pham S, Prjibelski AD, Pyshkin AV, Sirotkin AV, Vyahhi N, Tesler G, Alekseyev MA, Pevzner PA. 2012. SPAdes: a new genome assembly algorithm and its applications to single-cell sequencing. J Comput Biol 19:455–477.

103. Nadalin F, Vezzi F, Policriti A. 2012. Gapfiller: a *de Novo* assembly approach to fill the gap within paired reads. BMC Bioinformatics 13.

104. Sharma, A.K.; Dhasmana, N.; Dubey, N.; Kumar, N.; Gangwal, A.; Gupta, M.; Singh, Y. Bacterial Virulence Factors: Secreted for Survival. Indian J. Microbiol. 2017, 57, 1–10.

105. McClure, J.A.M.; Lakhundi, S.; Kashif, A.; Conly, J.M.; Zhang, K. Genomic Comparison of Highly Virulent, Moderately Virulent, and Avirulent Strains from a Genetically Closely-Related MRSA ST239 Sub-Lineage Provides Insights into Pathogenesis. Front. Microbiol. 2018, 9, 1531.

